# Coordinated active repression operates via transcription factor cooperativity and multiple inactive promoter states in a developing organism

**DOI:** 10.1101/2024.02.05.577724

**Authors:** Virginia Pimmett, Maria Douaihy, Louise Maillard, Antonio Trullo, Pablo Garcia Idieder, Melissa Costes, Jeremy Dufourt, Helene Lenden-Hasse, Ovidiu Radulescu, Mounia Lagha

## Abstract

Refining transcriptional levels via active repression in an euchromatic context represents a critical regulatory process. While the molecular players of active repression are well described, their dynamics remain obscure. Here, we used *snail* expression dynamics as a paradigm to uncover how repression, mediated by the Snail (Sna) repressor, can be imposed within a developing tissue. Combining live imaging and mathematical modeling, we show that Sna-mediated repression is cooperative and that cooperativity is primarily mediated by the distal enhancer. Repression shifts transcription bursting dynamics from a two-state ON/OFF regime to a three-state repressed regime with two temporally distinct OFF states. Mutating Sna binding sites suggests that repression introduces the long-lasting inactive state, which is stabilized by cooperativity. Our approach offers quantitative insights into the dynamics of repression and how transcription factor cooperativity coordinates cell fate decisions within a tissue.

## Introduction

Cell fate specification critically depends on differential gene expression. Cells are specified through the concomitant transcriptional activation of key lineage specifying factors and the repression of alternative fates. The coordination of gene activation is particularly important during the development of multicellular organisms where multipotent cells must choose distinct differentiation routes in an orchestrated manner. Thanks to functional genomics approaches, how the combinatorial action of transcription factor (TF) elicits the activation or repression of developmental promoters is relatively well understood. Downregulation of a gene is typically achieved by repressors, TFs that recruit co-repressors to reduce or silence gene expression. Depending on their range of action, they can be categorized into long or short-range repressors. Some (co-)repressors, such as Groucho/TLE, act over large distances and mediate long-range repression by silencing the entire locus. In contrast, short-range repressors function locally (50-150 bp) to inhibit the basal transcriptional machinery without interfering with more distant activators^1^. At the molecular level, several mechanisms have been proposed including direct competition between activators and repressors for a shared DNA binding site or ‘quenching’ of closely located activators and members of the basal transcription machinery^2^. A third and non-exclusive mechanism is the recruitment of histone deacetylases (HDACs), condensing chromatin and restricting access to the promoter^1^.

Two modes of transcriptional repression can be distinguished: the classical silencing in the context of heterochromatin or reduction in the context of a euchromatic environment, referred to as active repression. Contrary to Polycomb-mediated gene silencing, much less is known concerning active repression^3^. Yet, because of its fast establishment, reversibility, and capacity for partial reduction of expression, active repression stands as an optimal mode of gene expression control during periods of rapid decision-making. Failure of repression can lead to developmental defects and diseases such as cancer, as exemplified by the regulation of the Epithelial to Mesenchymal Transition (EMT). This fundamental cellular process is instructed by the conserved pro-EMT *snail* family, composed of Snail/Slug, Twist and ZEB1, acting as both activators and repressors^4^. Snail (Sna) is a zinc finger transcription factor primarily acting as a repressor, but reported to also act as an activator in some contexts^5^. Sna plays a critical role for correct completion of EMT, such as during *Drosophila* gastrulation or vertebrate neural tube formation^6^. Sna overexpression is sufficient to induce EMT^7^ and tumorigenesis^8^. Thus, tuning Sna levels and the network induced by this repressor TF is critical.

While repressor identities are well known, their impact on transcriptional kinetics is much less described. Transcription is inherently dynamic and occurs in pulses known as transcription bursts^9^. Transcription bursts are due to the stochastic switching of the promoter between permissive active states (ON, from which Pol II can initiate) and inactive states (OFF) of multiple timescales^9^. The mean RNA production depends on the switching rates between these states and on the active state production rate^10^. Repressors can in principle modulate any or all state switching rates, resulting in fine-tuning of repression rather than an all-or-nothing process. However, the scheme of state switching and the specific parameter(s) modulated by repression remain unknown.

In this study, we use *snail* expression dynamics as a paradigm to uncover how active repression, mediated by the Sna short-range repressor, can be imposed within a developing tissue. We take advantage of the power of quantitative live imaging to monitor endogenous *sna* transcription and protein dynamics in single nuclei prior to a major developmental decision, the EMT. Using novel theoretical approaches, we uncovered and quantified the timescales of the kinetic bottlenecks tuning transcription during repression. Based on experimental measurements, we propose a stochastic model of transcriptional dynamics and repression. In this model, repression adds a new long OFF state to the two-state unrepressed dynamic and modulates the stochastic switching rates cooperatively. Numerical simulations of this model suggest that the cooperativity between repressors contributes to coordination of repression within a tissue.

## Results

### Monitoring transcriptional repression in vivo

To decode the dynamics of transcriptional repression, we focused on a model gene, *snail* (*sna)*, which undergoes partial repression in the early blastoderm embryo. This gene encodes a key TF instructing the mesodermal fate and subsequent EMT prior to gastrulation. To examine the endogenous dynamics of *snail* expression in real time, we inserted a 24xMS2 array^11^ into the 3’ UTR of the endogenous *snail* gene using CRISPR-mediated recombination (*sna^MS^*^2^; **Figure 1A**). When both maternal and paternal alleles are tagged, the resulting *sna^MS^*^2^ flies are homozygous viable, and MS2 reporter expression matched the endogenous *sna* signals, as shown by single mRNA fluorescence *in situ* hybridization (smFISH) experiments **(Supplementary Figure 1A-B**).

**Figure 1:**
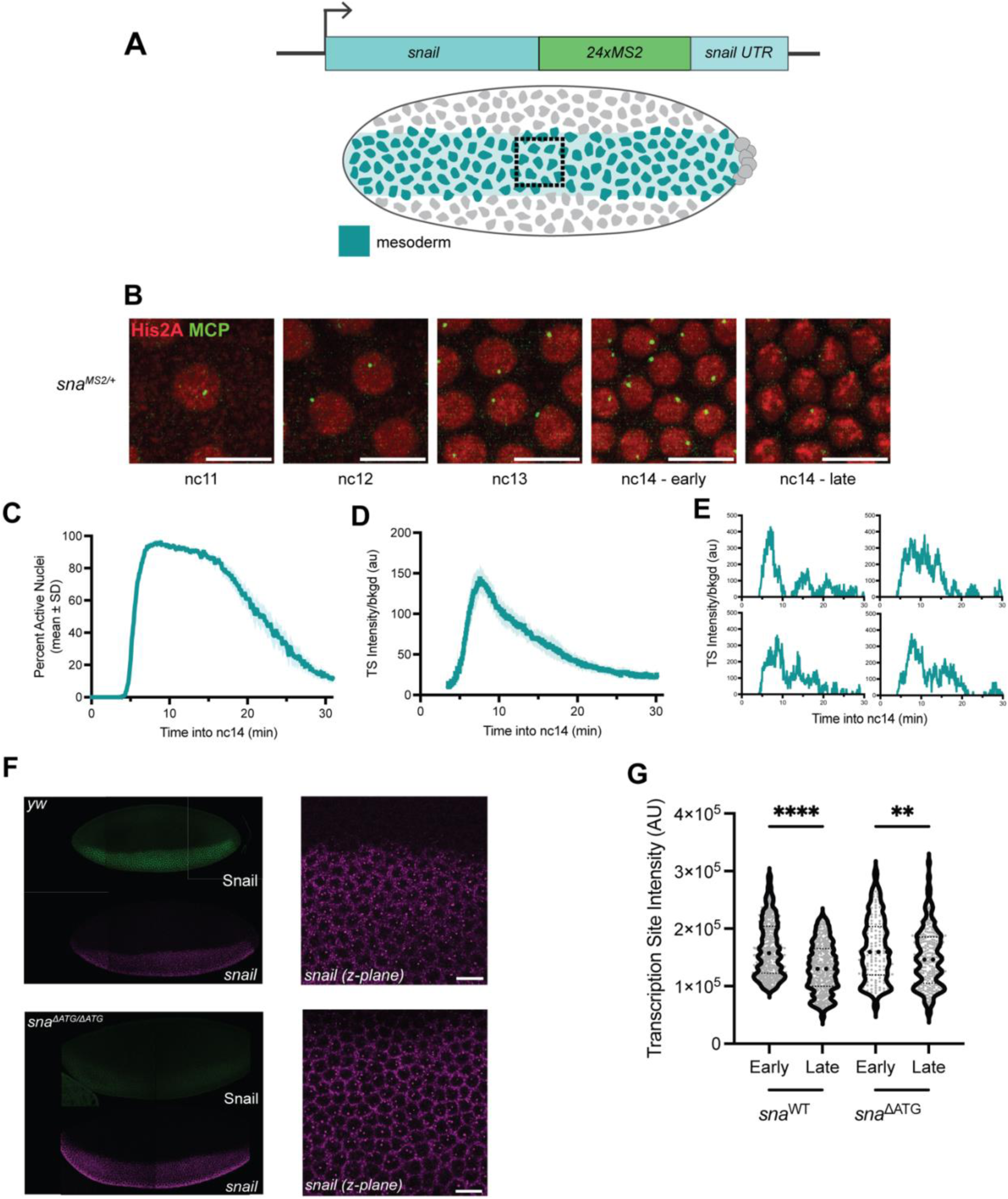
Live imaging of *snail* endogenous transcription demonstrates two regimes including a coordinated rapid transcriptional repression. A) Schematic view of *sna^MS2^* allele (above) and expression domain in the embryonic mesoderm (below, teal). The box indicates the restricted imaging area. B) Maximum intensity Z-projection of representative nuclei showing MS2/MCP-eGFP-bound puncta and nuclei (His2A-mRFP) in sequential nuclear cycles. Images were taken from a heterozygous embryo expressing *sna^MS2^* (**Supplementary Movie 1**). Scale bar represents 10 µm. C) Instantaneous activation percentage (mean ± SEM) curves of ventral nuclei during the first 30 min of nc14. Time zero is from anaphase during nc13-nc14 mitosis. D) Fluorescence intensity of actively transcribing nuclei (mean ± SEM) for *sna^MS2/+^* nuclei during the first 30 minutes of nc14. Time zero is from anaphase during nc13-nc14 mitosis. E) Sample single nucleus fluorescence traces in the first 30 minutes of nc14. Time zero is from anaphase during nc13-nc14 mitosis. F) *sna^ΔATG/ΔATG^* embryos demonstrate absence of Snail protein (green) and increased nascent transcription activity by smFISH relative to control embryos. Scale bar represents 10 μm. G) Quantification of endogenous *sna* and *sna^ΔATG^* transcription site intensity divided by background in early and late nuclear cycle 14 embryos via smFISH. Statistics: *sna^MS2/+^*: N=6 embryos, n=484 nuclei. smFISH: N=3 *sna^ΔATG/ΔATG^* embryos for early and late smFISH time points as determined by membrane invagination. Significance is indicated using a Kolmogorov- Smirnov test.

To image transcription, MS2 coat protein is fused to eGFP (MCP-eGFP) and provided maternally along with a fluorescently-tagged histone (His2A-mRFP). In combination with the paternally provided *sna^MS^*^2^ allele, transcription is visible as bright nuclear foci (**Supplementary Movie 1**). Signal intensity was retrieved in 3D and tracked through nc13 and nc14, using mitosis as a temporal ‘time-zero’ reference for each nuclear cycle.

We characterized the dynamics of endogenous *sna* transcription at the single nucleus level in embryos heterozygous for *sna^MS^*^2^. Transcription is detectable in living embryos as early as nc11 (**Figure 1B**). While transcription was stable throughout nc11-13, the dynamics in nc14 evolved significantly over time. Reactivation after mitosis in nc14 is rapid and synchronous but declines after a short plateau (**Figure 1C-D, Supplementary Figure 1C**). This change of regime and transcriptional attenuation is reflected in the individual nuclear traces as well, where TS intensity declined after peaking without completely vanishing (**Figure 1E, Supplementary Figure 1D**).

We turned to identifying a putative repressor of *sna* in nc14. As Sna is a transcription factor known to act as a repressor in *Drosophila* and to form an autoregulatory loop in other species^12,13^, we examined whether Sna formed an auto-repressive loop in the early embryo. To test Snail function while maintaining *snail* transcription as a readout, we created a protein-null allele (*sna^ΔATG^*) (**Figure 1F, Supplementary Figure 2A-D**). Using smFISH, we observed that the loss of Sna resulted in a derepression of *sna* transcription in late nc14 (**Figure 1G**). Thus, it appears that while *sna* transcriptional activity is stable during early embryogenesis, it undergoes rapid evolution in nc14 driven at least in part by Sna itself.

### Snail repression results in a non-stationary transcriptional regime

To access the kinetic parameters driving *sna* expression, we employed our previously developed deconvolution pipeline^14–16^ to extract the sequence of Pol II initiation events from the MS2-MCP-GFP signal for each single nucleus. Critically, this process does not rely on the arbitrary assignment of bursts to the signal and is valid for both stationary and non-stationary signals.

In brief, we consider the intensity trace of each spot to be a convolution of multiple concurrently transcribing polymerases and model the contribution of a single polymerase, assuming full processivity, constant speed and a negligible retention time at the transcription site (see **Methods**). To estimate the dwell time comprising Pol II elongation and transcript retention at the TS, we used signal autocorrelation^17^ (**Supplementary Figure 3A-B**); the resulting values are similar to those obtained from published Pol II speed measurements^18^. Using deconvolution, fluorescence traces can thus be converted to polymerase initiation events for single nuclei. Single nuclei can then be assessed as an ensemble of polymerase initiation events over time (**Figure 2A, B**).

**Figure 2:**
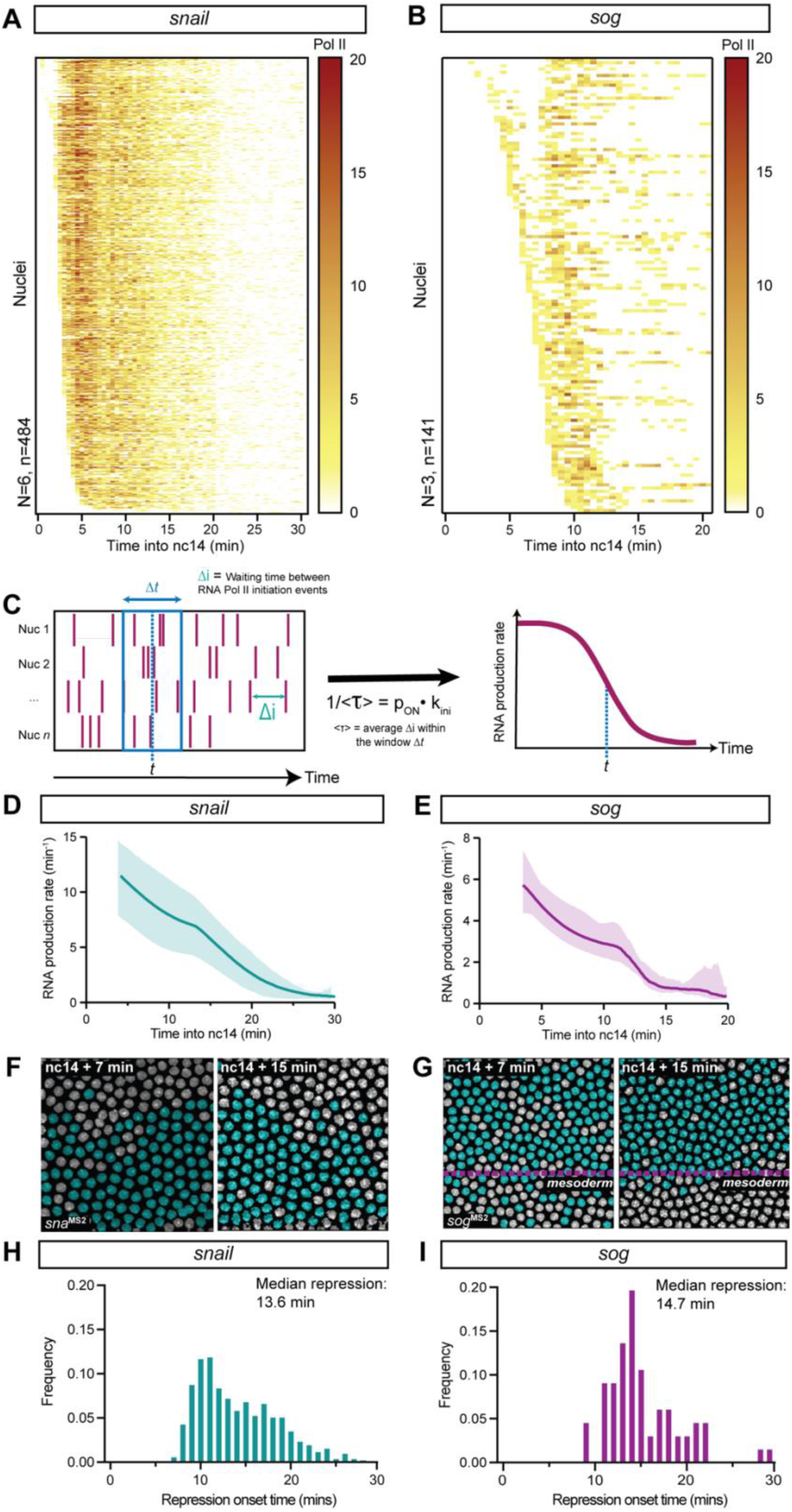
Deconvolution of transcription in living embryos reveals gene-specific behaviour in repression onset. A-B) Heatmap showing the number of polymerase initiation events in nc14 for *sna^MS2/+^* (A) and *sog^MS2/+^* (B) as a function of time. Each row represents one nucleus, and the number of Pol II initiation events per 30 s bin is indicated by the bin color. C) Deconvolution of the transcriptional site intensities into RNA polymerase II initiation events over time^18,19^. The average waiting time (*<τ>*) between polymerase initiation events is calculated for all nuclei within a sliding time window (*Δt*). The inverse of *<τ>* is the product of the probability to be active, denoted *p_ON_*, and the polymerase initiation rate (*k_ini_*) for the given time window. The inverse value is plotted over time as a proxy for the stability of the underlying transcriptional kinetic regime. Stationarity is denoted by a slope ≈ 0. D-E) Kinetic parameter stability as a function of time for *sna^MS2/+^* (D) and *sog^MS2/+^* (E) transcription expressed as the product of the probability to be active (*p_ON_*) and the RNA polymerase II initiation rate (*k_ini_*). F-G) False-coloured projections from live imaging of *sna^MS2^*^/+^ (F) and *sog^MS2^*^/+^ (G) embryos, with active nuclei indicated in teal and inactive in grey (**Supplementary Movies 1,2**). H-I) Distribution of switching times for initiation of stable repression in nuclear cycle 14 for *sna^MS2^* (H) and *sog^MS2^* (I) as determined using Bayesian Change Point Detection. Statistics : *sna^MS2/+^* : N=6 embryos, n=448 nuclei. *sog^MS2/+^* : N=3 embryos, n=141 nuclei.

We extracted the waiting times between polymerase initiation events for each nucleus and quantified the mean waiting time between polymerase initiation events (*<τ>*) in a sliding window (**Figure 2C**). *<τ>* summarizes the repression kinetics, representing the inverse mean RNA production, and is thus directly related to the product of *p_ON_*, or the probability to be in a productive state, and *k_ini_*, or the initiation rate while in the productive state^19^. The temporal profile of *p_ON_ k_ini_* = 1/*<τ>* is a readout of the stability of transcription. Using a sliding window, the temporal evolution of *p_ON_ k_ini_* (or RNA production rate) can be tracked across time. We examined the temporal evolution of the RNA production rate for *sna* in nc14 (**Figure 2D**) and compared it to another target of Snail-mediated repression, *short gastrulation* (*sog*, **Figure 2E, Supplementary Movie 2**)^20^. Unlike *sna*, which undergoes a partial repression, *sog* is fully silenced by the action of Sna^21^ (**Figure 2F, G**). For both *sna* and *sog*, the mean waiting time between polymerase initiation events was non-stationary across nc14 (**Figure 2D, E**). Thus, stationary repression is progressively imposed regardless of whether transcription was attenuated in nc14 (*sna*) or completely repressed (*sog*).

As the population-level signal is non-stationary, we employed the Bayesian change point detection (BCPD) algorithm^22^ to identify the timing of change between a transcriptionally unrepressed and repressed program at the single nucleus level (see **Methods**). Importantly, this algorithm identified the point at which the repressive regime is stabilized (**Supplementary Figure 4A-D**). Interestingly, the inter- nuclear coordination of repression, represented by the breadth of repression onset times, was weaker for incomplete repression (*sna*, **Figure 2H**) compared to complete repression (*sog*, **Figure 2I**). We also examined whether repression was implemented similarly between both alleles of *sna* by analyzing *sna^MS2^/sna^MS2^* embryos. The median time where repression achieved stationarity for the first- and second- activated allele was similar to both each other and a randomly selected pool of alleles (**Supplementary Figure 5A-F**), indicating repression is implemented uniformly across alleles. We conclude that, although repression is applied progressively, stationary transcription is reached after a certain onset time, and variations in this time can be used to quantify the coordination of repression.

### Snail-mediated active repression involves Sna cooperative action

To examine the interplay between endogenous Sna protein and its target genes *sna* and *sog*, we employed the LlamaTag system^23^ and created a CRISPR *snail^Llama^* (**Figure 3A, B**). The LlamaTag system relies on a nanobody, targeting a fluorescent protein, fused to a protein of interest and the presence of a free fluorescent detector. Once Sna protein is translated in the cytoplasm, the Llama nanobody binds to free maternally-deposited GFP and is then imported in the nucleus. The subsequent increase in nuclear fluorescence signal provides a readout of Sna nuclear levels.

**Figure 3:**
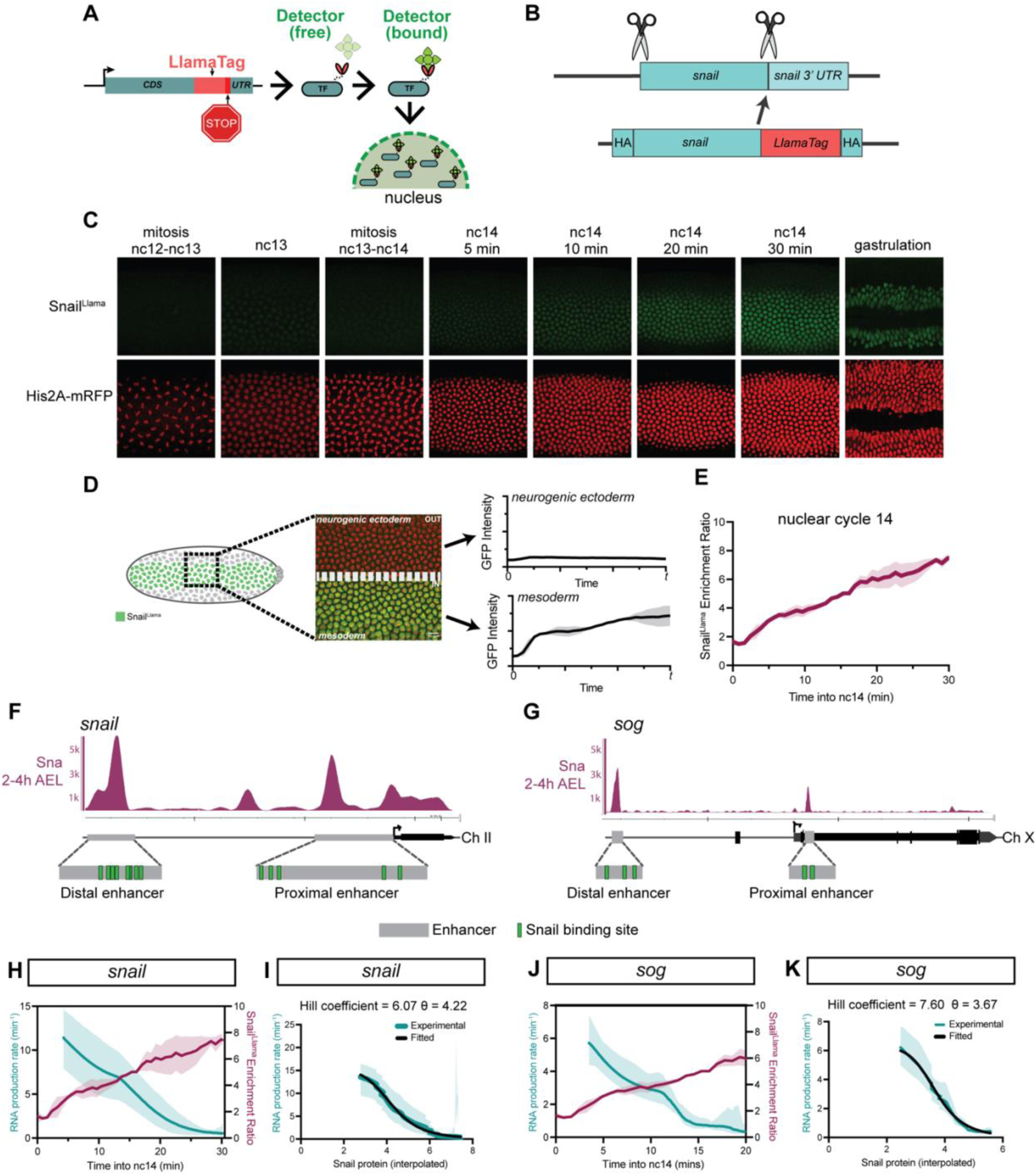
Capturing Snail protein dynamics and its correlation with transcription repression in living embryos. A) Schematic demonstrating the principle of the LlamaTag system. B) Schematic indicating CRISPR-mediated genome editing of the endogenous *sna* locus to introduce the LlamaTag nanobody. C) Representative maximum intensity Z-projections of Sna^Llama^ nuclear signal (above) and nuclei (His2A-mRFP, below) from nuclear cycle 12-13 mitosis until gastrulation. See also **Supplementary Movie 3**. D) Schematic of Sna^Llama^ analysis showing the imaging window on the embryo, with the mesoderm/neurogenic ectoderm boundary indicated. The nuclear GFP intensity for both the mesoderm and neurogenic ectoderm was quantified as a function of time. E) Nuclear Sna^Llama^ enrichment ratio in the first 30 minutes of nc14 calculated as the mean nuclear GFP signal in the mesoderm relative to the neurogenic ectoderm, expressed as mean ± SD. F-G) Loci and embryonic enhancer sequences (grey) of *sna* and *sog*, with Sna binding sites (green) indicated. H,J) *p_ON_·k_ini_* (mean RNA production) expressed as a function of time for *sna^MS2^* (H, teal) and *sog^MS2^* (J, purple) and Snail protein enrichment ratio (red) in nc14. The mean RNA production expressed as mean ± SD. I,K) Hill fitting for *sna^MS2^* (I) and *sog^MS2^* (K) with the Hill coefficient and θ indicated above. θ represents the repressor concentration reducing transcription intensity to half. The uncertainty interval of the Hill coefficient was computed as the standard deviation of the Hill coefficient coming from the fitting of different *sna^MS2^* movies. Coloured line indicates experimental kinetic parameter stability as a function of time expressed as the product of the probability to be active (*p_ON_*) and the RNA polymerase II initiation rate (*k_ini_*) in nc14 (mean ± upper/lower bounds). Black line indicates Hill function fit. Statistics: *sna^MS2/^*^+^: N=6 embryos, n=448 nuclei; *Sna^Llama/+^*: N=3 embryos; *sog^MS2/+^*: N=3 embryos, n=141 nuclei.

The *sna^Llama^* CRISPR allele allowed us to quantify with high spatio-temporal resolution the levels of endogenous Sna protein (**Figure 3C, E-F**) by tracking the nuclear GFP signal as a proxy for Sna concentration in the nucleus marked by His2A-RFP. Despite the early accumulation of *sna* transcripts (**Figure 1B**), nuclear accumulation of Sna protein was detected weakly in nc13 (**Figure 3C, Supplementary Movie 3**). By comparing signal within the mesoderm to that in a neighboring tissue where *sna* is not expressed (neurogenic ectoderm, **Figure 3D**), we observed Sna nuclear protein levels in nc14 continuously increase in the presumptive mesoderm (**Figure 3E**).

Sna has previously been demonstrated to bind known enhancers of both *sna* itself as well as *sog*^24^ (**Figure 3F, G**). Consistent with the idea that Sna acts as a transcriptional repressor, we observed an anticorrelation between Sna protein levels and RNA production rates of both *sna* and *sog* (**Figure 3H, J**). We next wanted to quantitatively characterize their relationship. The input/output relationship is steep for both *sna* and *sog*, indicating thresholding effects. Such relationships are typically fitted with a Hill equation^25^ that accesses a key parameter, the Hill coefficient *n* (representing the degree of cooperativity). The Hill coefficient for *sna* during nc14 was 6.07 (**Figure 3I**). Similarly, the relationship between input Sna protein and output *sog* transcription also indicates high-degree cooperativity (Hill coefficient 7.6, **Figure 3K**).

Collectively, the relatively high Hill coefficients indicate that Sna protein may act cooperatively to elicit repression via its own *cis-*regulatory regions and through *sog* enhancers. We note however, that the Hill equation model is purely phenomenological and does not account for the details of the underlying mechanism.

### Sna cooperativity is differentially mediated through distinct cis-regulatory regions

We exploited previously characterized *sna* BAC reporter lines^26^ to explore the role of cooperativity underlying Snail-mediated repression. *Sna* is regulated by a pair of non-redundant enhancers, one proximal and one distal (**Figure 4A**). We performed quantitative imaging of the *sna* wild-type and enhancer deletion BACs in the mesoderm (**Figure 4B**) and characterized the relationship between RNA production rate and the Snail endogenous protein using a Hill function. Compared to the control, loss of the distal (i.e. shadow^27^, **Figure 4C,F**) enhancer (*sna^ΔDIST^*, **Figure 4D**) resulted in a complete loss of Sna cooperativity (Hill coefficient ≈1, **Figure 4G**) while the loss of the proximal enhancer (*sna^ΔPROX^*, **Figure 4E**) demonstrated increased Sna cooperativity (Hill coefficient ≈4) (**Figure 4H**). Thus, it appears that Sna cooperative action is primarily decoded by the distal enhancer.

**Figure 4:**
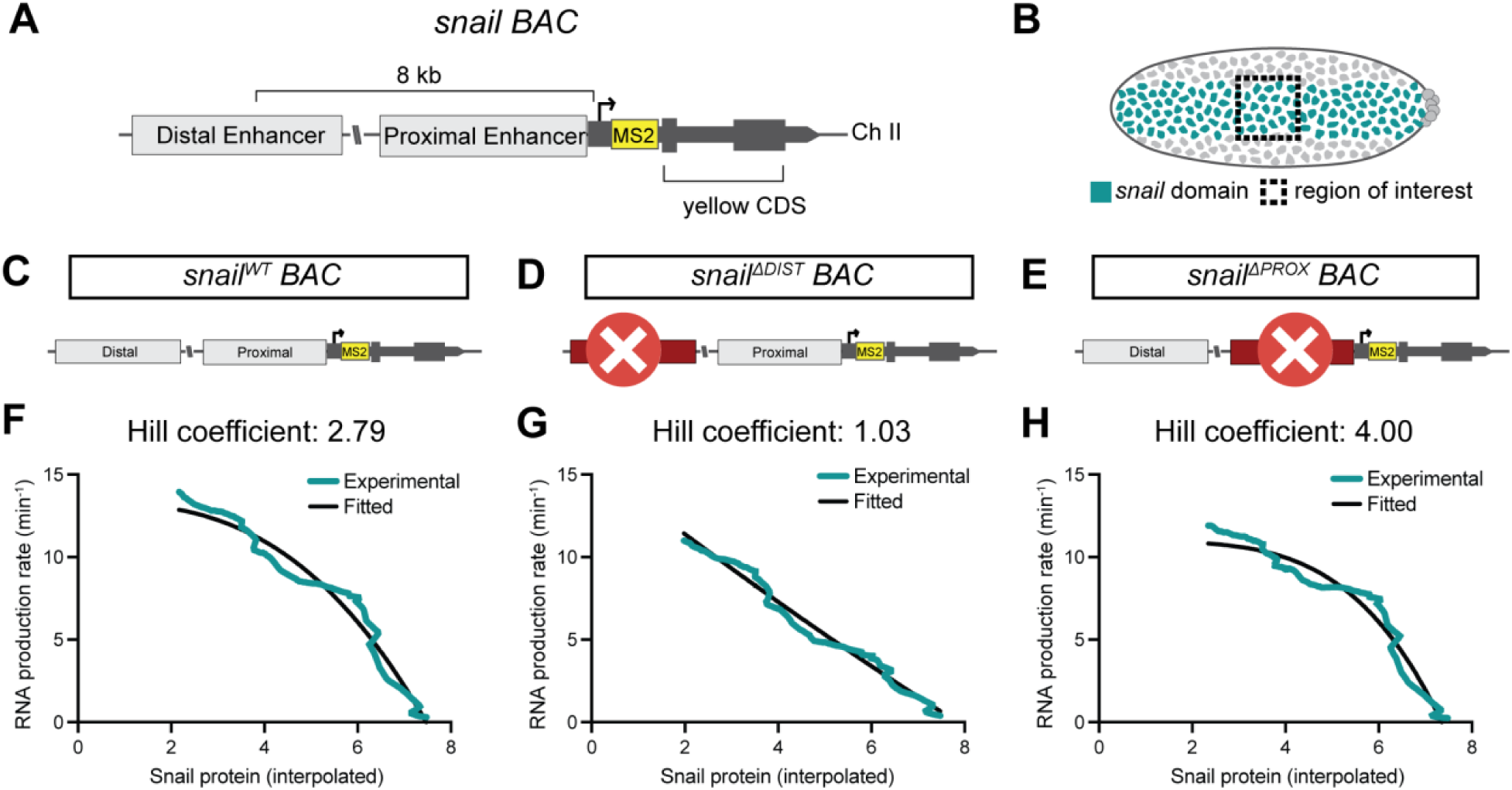
Sna cooperativity is differentially mediated by its proximal and distal enhancers. A-B) Schematic of *snail* BAC reporter construct (A) and expression domain (B, teal) with region of interest indicated (dashed box). C-E) Schematic of reporter constructs for *sna^WT^* (C), *sna^ΔDIST^* (D) and *sna^ΔPROX^* (E). F-H) Hill fitting for *sna^WT^* (F), *sna^ΔDIST^* (G) and *sna^ΔPROX^* (H) with the removed enhancer and Hill coefficient indicated. Coloured line indicates experimental kinetic parameter stability as a function of time expressed as the product of the probability to be active (*p_ON_*) and the RNA polymerase II initiation rate (*k_ini_*) in nc14. Black line indicates Hill function fit. Statistics: *snail^WT/+^* BAC N=4 embryos, n=341 nuclei; *snail^ΔDIST/+^* BAC N=5 embryos, n=344 nuclei; *snail^ΔPROX/+^* BAC N=3 embryos, n=217 embryos.

### Sna repressor acts by introducing a second non-productive state and modulating the promoter’s state switching rates

It remains an open question which aspect of the transcriptional kinetic cycle repression acts on. As the Hill equation is phenomenological in nature, we turned to transgenic reporter assays to establish a causal link between Snail binding and transcriptional repression, and to investigate the link between promoter dynamics and the imposition of repression.

The *sna* distal enhancer has a cluster of nine Sna binding sites (**Figure 5A**, pink bars). We employed our previously established transgenic reporter platform^21^ to investigate the effect of these Sna TF binding sites on transcription dynamics. We generated a series of transgenes where transcription is controlled by 4 types of enhancers (**Figure 5B-E)**: the wild type *sna* distal enhancer (**Figure 5B**, *snail^Distal^*), a mutated distal enhancer with every other (ie. 5/9) Sna binding site removed (**Figure 5C**, *snail^DistalAlt^*), with 8/9 Sna binding sites removed (**Figure 5D**, *snail^DistalMut^*), or a fragment of the distal enhancer with no Sna sites (**Figure 5E**, *snail^DistalCore^*). Following deconvolution, we observed the *snail^Distal^* transgene had globally lower numbers of polymerase initiation events compared to *snail^DistalAlt^*, *snail^DistalMut^* and *snail^DistalCore^* (**Figure 5F-I**). Thus, the *sna* distal enhancer is derepressed in the absence of Sna binding sites.

**Figure 5:**
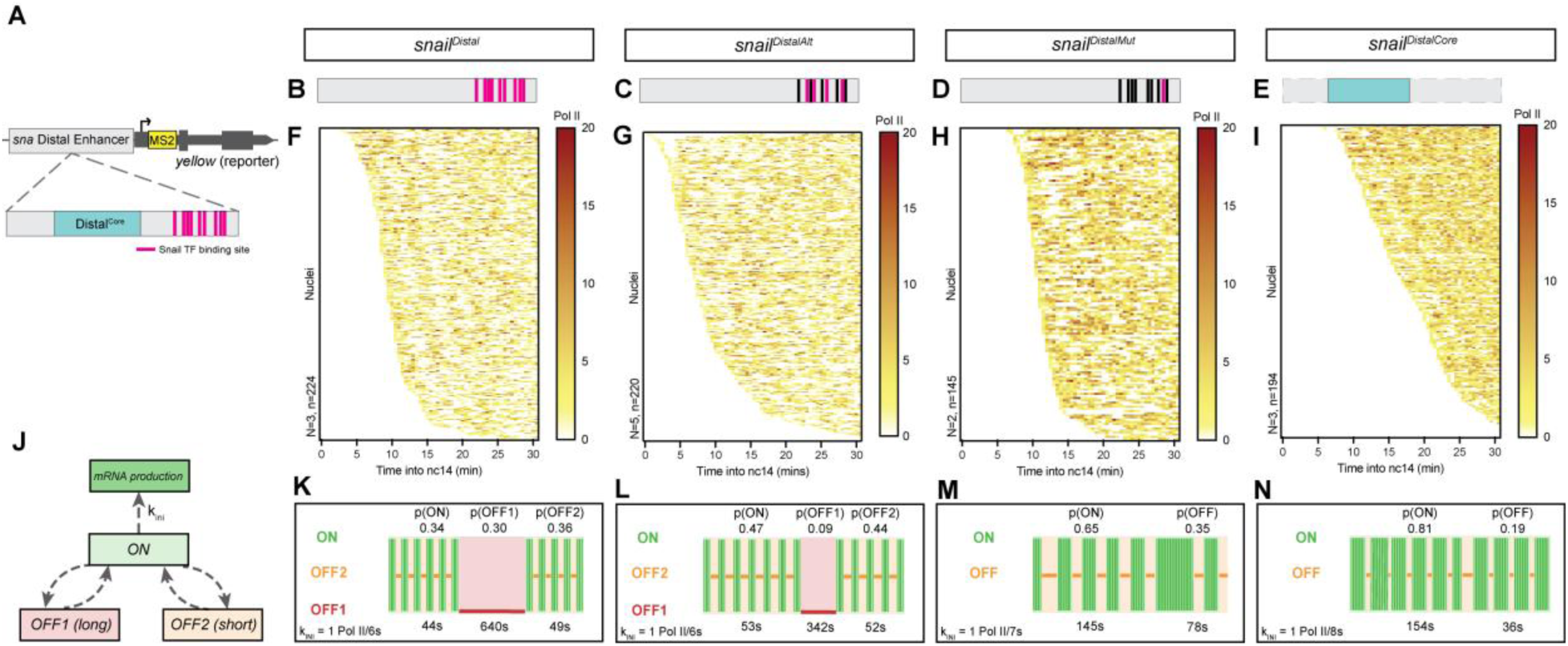
Snail repressor introduces a second non-productive state and modulates promoter state switching rates. A) Schematic of the *sna* distal enhancer transgene with the ‘core’ enhancer module^57^ (blue) and Snail binding sites (pink) indicated. B-E) Enhancer transgene schematics for the *snail^Distal^* enhancer (B), *snail^DistalAlt^* with mutated Sna binding sites (C), *snail^DistalMut^* with mutated Sna binding sites (D) and *snail^DistalCore^* with the ‘core’ enhancer module only (E). F-I) Heatmap showing the number of polymerase initiation events in nc14 for *snail^Distal^* (F), *snail^DistalAlt^* (G), *snail^DistalMut^* (H) and *snail^DistalCore^* (I) as a function of time. Each row represents one nucleus, and the number of Pol II initiation events per 30 s bin is indicated by the bin color. J) Topology of the three-state kinetic model with non-sequential OFF1 and OFF2 states. K-N) Representation of estimated bursting dynamics for *snail^Distal^* (K), *snail^DistalAlt^* (L), *snail^DistalMut^* (M) and *snail^DistalCore^* (N). Permissive ON state durations are depicted in green and inactive OFF states in red and orange, and probabilities of each state shown above. Statistics: *snail^Distal^* N=3 embryos, n=224 nuclei; *snail^DistalAlt^* N=5 embryos, n=220 embryos; *snail^DistalMut^* N=2 embryos, n=145 nuclei; *snail^DistalCore^* N=3 embryos, N=194 nuclei. See **Supplementary Movies 4-7**.

To further investigate the underlying kinetics of transcription driven by each enhancer, we examined the distribution of waiting times between polymerase initiation events. Briefly, this distribution can be fitted with a multi-exponential function that provides insight into the number of promoter states as well as their duration (T_State_) and probabilities (p_State_), and the polymerase initiation rate (k_INI_)^14^ (**Supplementary Table 1**).

The distribution of Pol II waiting times from the *snail^Distal^* construct could not be fitted with a bi-exponential function, meaning a two-state ‘random telegraph’ model was insufficient to describe the promoter states (**Supplementary Figure 6A-B**). A three-exponential fitting was sufficient, corresponding to a three-state model (**Figure 5J**) with one productive (ON) state and two non-productive states on the order of minutes (OFF1) or seconds (OFF2) of approximately equal probability (**Figure 5K**). The *snail^DistalAlt^* transgene also required a three-state model fitting, but interestingly showed a substantial decrease in the length and probability of the OFF1 state, with a higher probability of the ON state to compensate (**Figure 5L, Supplementary Figure 6C-D**). Transcription from the *snail^DistaMut^* enhancer, with 8/9 Sna binding sites mutated, was adequately fit by a two-state model (**Figure 5M, Supplementary Figure 6E**). Importantly, *snail^DistaMut^* transcription dynamics exhibit a loss of the long OFF1 state, at the expense of an increase in both the duration and probability of the productive ON state as well as a small increase in the duration of the single non-productive OFF state. This bursting regime was recapitulated in the *snail^DistaCore^* reporter, which was also well described by a two-state model with a longer and highly probable ON state and a single short non-productive state (**Figure 5N, Supplementary Figure 6F**). Interestingly, the initiation rate was consistent between all three reporter constructs, indicating that Sna binding does not affect Pol II firing from the ON state. Thus, the Sna repressor likely acts by introducing a new kinetic bottleneck, leading to a second non-productive state at the expense of the productive ON state and the pre-existing non-productive OFF state, and without affecting the polymerase initiation rate.

Multiple kinetic state topologies have been proposed for higher-order kinetic models^31^ with multiple productive and non-productive states. Based on our previous work, and in agreement with the deconvolution findings, we favour a non-sequential three state model of promoter dynamics (**Figure 5J**) where there is no direct transition between the temporally distinct long OFF1 and short OFF2 state. In this model, the transitions from ON to OFF1 and from ON to OFF2 are independent. A sequential model, as also described in the literature^15,16,28^, would require the promoter to systematically switch to OFF2 before transitioning to OFF1. We have shown that both the sequential and non-sequential models are compatible with the MS2 data^15,19^; however, the strong correlation between the two OFF states implied by the sequential model would be difficult to justify mechanistically.

We also conclude that one of the two OFF states of the three-state model is suppressed when transcription is driven by the *snail^DistaMut^* and *snail^DistaCore^* enhancers. In the *snail^DistalAlt^* construction the long OFF1 state occurs rarely (*p*_OFF1_=0.09 compared to *p*_OFF1_=0.3 in *snail^Distal^*), suggesting repression introduces the long OFF1 state. This conclusion is further supported by the distinct orders of magnitude of the OFF state durations: *T*_OFF1_ is on the order of seconds, and *T*_OFF2_ on the order of minutes in the three-state constructions, whereas the single *T*_OFF_ in the two-state constructions is clearly closer to the *T*_OFF2_ values.

In summary, we find that the Sna repressor acts by introducing a second long non-productive state and modulating all state switching rates, but it does not affect the polymerase initiation rate in the permissive state.

### The endogenous repressed state recapitulates transgenic reporter activity

Transgenic reporter experiments suggest that transcription bursting dynamics follow a three-state regime under repression, featuring an additional long OFF state that is not present in the absence of repression.

We hypothesized the autorepression of *sna* would also require a three-state topology, since there are several clustered Sna binding sites in the endogenous regulatory scheme (**Figure 3F**). We used the BCPD procedure previously outlined to isolate the stably repressed phase of *sna^MS2^* transcription (**Supplementary Figure 7A-B**) and fit it using various multi-exponential models. We found that a three-state model was required to capture the kinetics of endogenous *sna^MS2^* during stable repression (**Figure 6A-B, Supplementary Figure 7C-D**). This contrasted the unrepressed nc13, where a bi-exponential fitting could recapitulate the data (**Supplementary Figure 8A-F**).

**Figure 6:**
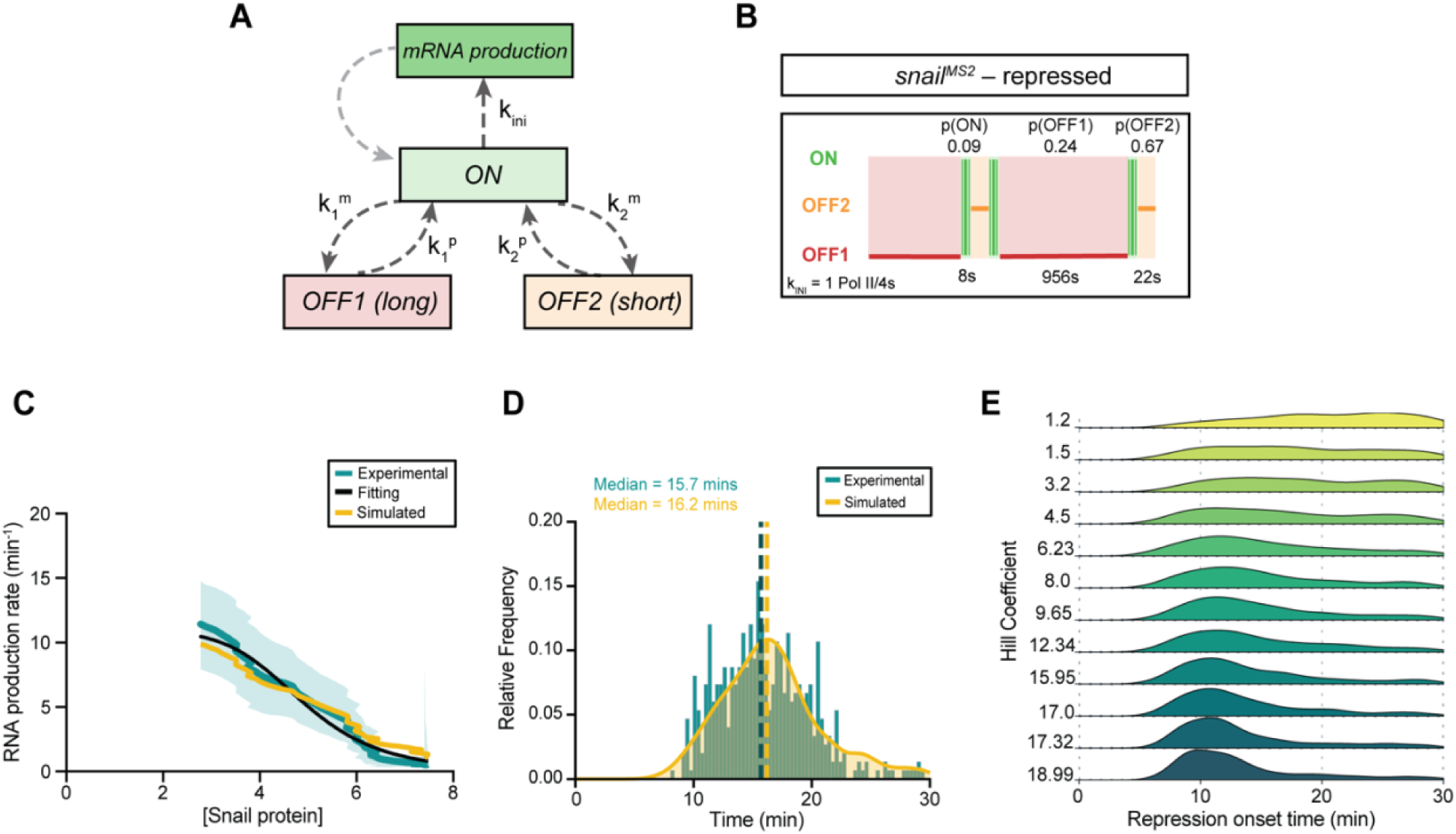
A minimal stochastic model reveals the impact of cooperativity on transcriptional repression. A) Scheme of the Markovian model. B) Representation of estimated bursting dynamics for *sna^MS2^* during stable repression in nc14 (**Supplementary Movie 1**). Permissive ON state durations are depicted in green and inactive OFF states in red and orange, and probabilities of each state shown above. C) Best stochastic model fit (yellow) parallel the Hill function model fit of modelled RNA production rate (black) and the experimentally derived values (teal) over 30 minutes of minimal stochastic model fitted transcription. D) Comparison of the BCPD-derived repression onset times for the experimentally-derived data (teal) and kernel density estimate for simulated data from C (yellow). Median repression onset time is indicated by dashed lines. E) Distribution of simulated repression onset time for various Hill coefficients.

Previous research has established a role for promoter-proximal polymerase pausing in the introduction of a novel promoter state^16^. We thus tested whether pausing was responsible for the introduction of a new rate-limiting step in transcription, by perturbing the levels of key pausing factors, Paf1^29^, a subunit of the NELF complex^30^ or Cyclin T^31^, required for pause release. We employed two independent RNAi-mediated knockdowns of *paf1*, RNAi-mediated knockdown of *Nelf-A*, as well as overexpression of *Cyclin T* in the early embryo. None of these perturbations altered the number of rate-limiting steps during stable *sna* repression (**Supplementary Figure 9A-O**).

Collectively these results suggest that under repression, the endogenous promoter switches between three temporally distinct states: a competent ON state from which Pol II initiates at a given rate, and two inactive states, one short at seconds-scale and a longer minutes-scale state. Because the three-state promoter topology persists in conditions where pause release is favored, we conclude that the extra rate-limiting step present during repression cannot be attributed to a paused state.

### A minimal stochastic model links cooperativity and coordination of repression

To gain quantitative insights into the relationship between Snail-mediated repression and transcriptional bursting, we developed a minimal stochastic model of transcriptional bursting. The model was designed to describe the active (nc13), the repression buildup (early nc14), and repressed regimes (mid-nc14) of *sna* transcription. We therefore used a three-state promoter model comprising two transcriptionally inactive states, a long one (OFF1) and a short one (OFF2), and a permissive (ON) state (**Figure 6A, Supplementary Figure 10A-B**).

To develop a minimal stochastic model, we first ensured that the state OFF1 is not accessible from the permissive ON during the active phase, corresponding to low concentrations of the putative repressor Sna. To achieve this, we considered that the parameter *k ^m^* (transition from ON to OFF1) decreases to zero when the concentration of Sna protein ([Sna]) approaches zero. Thus, when [Sna] is low, OFF1 can no longer be reached from ON, rendering the three-state model equivalent to a two-state model.

The parameters *k_2_* and *k_2_* (transition rate from ON to OFF2 and from OFF2 to ON) have smaller values in nc13 relative to nc14 (**Supplementary Table 1**) and were considered to increase with [Sna]. The parameter *k_1_^p^* (OFF1 to ON transition) is not available for nc13 because OFF1 is absent in the two-state model. However, it is available for nc14 and for the transcription regimes of sna enhancer reporter transgenes (**Figure 5**). When these reporters are arranged in order of decreasing repression 5 **Supplementary Figure 10C-G**), the value of *k_1_* increases, which leads us to propose a decrease in *k_1_* with increasing [Sna]. The limiting ([Sna] -> 0) value of *k_1_* for small concentration of [Sna] is a free parameter that is fitted from data. All the dependencies of state switching rates on [Sna] were modelled as Hill functions to account for cooperativity. Although there may be relationships between the values of these rates and the Hill coefficient (for instance, a large Hill coefficient may correlate with longer-lived states), to minimize the number of parameters and avoid overfitting, we considered a single coefficient for all the states and transitions. Finally, we considered that the polymerase initiation rate (k_ini_) does not depend on [Sna].

After implementing the active nc13 and repressed nc14 phase parameters (see **Methods**), our model remained with only three free parameters: the limiting ([Sna] -> 0) *k_1_^p^* value, the Hill coefficient *n* and the Sna repression threshold concentration θ. Gillespie simulations were used to generate synthetic nascent RNA data for this model and the three free parameters were fitted using the experimental RNA production rate data in nc14 (**Figure 6C**).

Next, we simulated the distribution of the repression onset time within the mesodermal population utilizing the fitted parameters (**Figure 6D**). The good agreement between the predicted and experimental repression onset time distributions validates the model.

The minimal stochastic model was then used to investigate the coordination of repression within a tissue. We simulated the model for many values of the Hill coefficient (corresponding to a spectrum of TF cooperativity) and Sna concentration threshold and computed the repression onset time distributions (**Figure 6E**). The width of this distribution, quantitatively defined as the interquartile range, serves as a measure of repression coordination. In general, for a fixed concentration of repressor, the distribution of the repression onset time narrows as the Hill coefficient increases (**Figure 6E**). Because a larger Hill coefficient indicates a higher degree of cooperativity, this suggests that Sna cooperativity contributes to the coordination of repression between nuclei.

## Discussion

Understanding the mechanisms by which gene transcription is dynamically attenuated within a developing tissue is a fundamental question. Here we use live imaging and mathematical modeling to quantitatively address this question in the context of *Drosophila* early development. We focus on *sna* as a model gene to extract the kinetics of promoter-switching during active repression, as well as the dynamics of the short-range repressor protein it encodes. By monitoring nascent mRNA and protein levels from endogenous loci in live embryos, we unveil 2 main features of active repression: (1) active repression introduces a new long OFF state to the two-state unrepressed dynamics; (2) repression modulates the stochastic switching rates cooperatively. Furthermore, we predict that the inter-nuclear coordination of repression is augmented by a high degree of repressor cooperativity.

### Transcription kinetics during active repression

The analysis of the distribution of waiting times between polymerase during the repression phase revealed the existence of two distinct OFF periods, one in the range of seconds and a prolonged one in the range of minutes. The molecular nature of these rate-limiting steps can only be an interpretation. Because of the comparison between varying number of Sna binding sites (**Figure 5**), we propose that the long OFF1 state, apparent only in repression, corresponds to a repressor-bound state. While the residence time of specific repressors has yet to be detailed *in vivo*, it is well-demonstrated that, apart from a few exceptions, activating TFs remain bound to DNA for up to tens of seconds^32^ in both *Drosophila* and vertebrates. How can we reconcile typical TF residence time with the prolonged OFF1 state resulting from Sna binding? Given the dependency between OFF1 duration and the number of Sna binding sites and their arrangement, we propose Sna binds in a cooperative manner. Cooperative binding of Sna repressors to DNA may stabilize a longer OFF1 state. A similar scenario of a prolonged promoter state driven by TF cooperativity has recently been proposed in yeast^33^. Albeit reported for activation and not repression, the rationale is nonetheless similar: TF exchange, possibly via cooperative binding, can increase the duration of a rate-limiting step during transcription.

### Snail-mediated repression: a cooperative action

The cooperativity of repression manifests as a steep response of RNA production to protein concentration. Our mathematical model suggests that the high degree of cooperativity governing Sna-mediated repression might optimize inter-nuclear coordination of repression within a tissue.

It is well-demonstrated that TFs bind DNA cooperatively^34,35^. Multiple examples point to the cooperative action of a combinatorial set of TFs to activate or silence cis-regulatory elements, but the underlying mechanisms remain unclear. Here, by directly measuring the relationship between nuclear protein concentration and transcriptional output in live embryos, we provide a quantitative estimation of cooperativity. By examining two transcriptional targets, *sna* and *sog*, expressed in the same tissue at a similar developmental state, we obtained an estimation of Sna-mediated cooperativity, with Hill coefficients in the range of 6 and 7.6 for *sna* and *sog* respectively. This is similar to that measured for the repressor Knirps in *Drosophila*^36^.

Cooperativity can occur through protein-protein and protein-DNA interactions, which imposes a particular arrangement of TF binding sites, but can also be achieved with more flexible arrangements such as a local change in DNA structure^37^ or mediated through competition with nucleosomes^38^. In principle, all these modes of cooperativity (DNA mediated, TF-TF interaction, or nucleosome-mediated) can lead to a high Hill coefficient. Although both *sna* enhancers contain Sna binding sites, they are clustered in the distal enhancer while their arrangement is more flexible in the proximal (**Figure 3**). Interestingly, mutation analyses suggest that these so-called ‘redundant’ enhancers decode Sna repressor action very distinctly, supporting increased cooperative action at the distal enhancer compared to the proximal.

Future investigations, including promising single molecule technologies as single molecule footprinting assays^39^ coupled to theoretical models^40^, would be required to elucidate which TF co-occupy the same enhancer DNA molecule *in vivo*. Such TF co-occupancy mapping would be greatly enhanced by the quantification of TF binding kinetics. Recent advances in single molecule imaging, including the exciting possibility of imaging temporally-evolving repressor ‘hubs’^41,42^, promise encouraging future insights.

In summary, by monitoring transcription and nuclear transcription factor levels in a developing embryo, we have uncovered kinetic bottlenecks governing repression. Our findings support a multiscale bursting model characterized by both short and long transcriptionally inactive periods. In this model, the initiation of repression results in prolonged non-productive periods governed by slow timescales. Looking ahead, we expect that the framework of analysis and results of this study will set a foundation for understanding repression dynamics in these more complex vertebrate models of development.

## Data availability

BurstDECONV source code is available through GitHub at https://github.com/oradules/BurstDECONV, and also on Zenodo at https://zenodo.org/record/7443044. SegmentTrack is available via GitHub at https://github.com/ant-trullo/SegmentTrack_v4.0) and the post-processing tool developed for this study at https://github.com/ant-trullo/SpotsFiltersTool. NucleiTracker3D is available via GitHub at https://github.com/ant-trullo/NucleiTracker3D. The BCPD algorithm is available via GitHub at https://github.com/mariadouaihy/BCPD_inhomogeneous_transcriptional_signal. Raw imaging data is available upon request.

## Acknowledgements

We are grateful for Mattias Mannervik for sharing the *UAS:CyclinT* fly stock, and Tamar Juven-Gershon and Anna Sloutskin for sharing the *w^coffee^* repair matrix, *vasa-*Cas9 stock, and assistance with the ssODN co-CRISPR system.

We thank Mattias Mannervik, Jean-Christophe Andrau, Arnaud Krebs, Marcelo Nollmann, John Reinitz and members of the Lagha lab for their critical reading of the manuscript. We thank John Reinitz and Edouard Bertrand for insightful discussions. We are grateful to Amandine Palandri for help with fly handling. We thank Rachel Topno for assistance with autocorrelation-based dwell time estimates. We acknowledge the Montpellier Ressources Imagerie facility (France-BioImaging) and the Biocampus Drosophila facility of Montpellier.

This work was supported by the ERC SyncDev starting grant to M.L. and the ANR HubDyn grant to M.L. M.D. is supported by the CNRS and University of Chicago Joint PhD programme. L.M. is supported by a fellowship from the LabMuse/University of Montpellier PhD program and a PhD fellowship from La Ligue Contre Le Cancer. M.L., O.R., and A.T. are sponsored by CNRS.

## Contributions

Conceptualization: ML, OR, VP, MD

Investigation/Data acquisition: VP, LM, MD, MC, PGI

Image analysis : AT,VP,LM

Mathematical modeling and code development : MD, OR

Stochastic model implementation : OR

Design and supervision of *sna^Llama^* plasmid construct cloning: JD

Cloning of *sna^Llama^* plasmid construct : HL-H

Funding acquisition: ML, OR

Project administration: ML

Supervision: ML, OR

Writing original draft: VP, ML

Writing, review and editing: OR, MD, LM, AT

## Methods

### Fly Husbandry

All crosses were maintained at 25°C. Transgenic and CRISPR lines were maintained as homozygous stocks unless otherwise noted (**Supplementary Table 2**). For live imaging of single MS2 allele crosses, homozygous males carrying the allele of interest were crossed with homozygous females bearing a *nos>MCP-eGFP-His2Av-mRFP* transgene. For live imaging of *snail^Distal^-*related transgenes, females heterozygous for *nos>MCP-eGFP-His2Av-mRFP* were crossed to males homozygous for the transgene of interest. For live imaging of RNAi and overexpression experiments, homozygous males carrying *mat- α>gal4 ; nos>gal4, nos>MCP-eGFP-His2Av-mRFP* were crossed with homozygous females bearing the RNAi or overexpression transgene of interest. F1 virgin females were then crossed to males bearing the *sna*^MS2^ allele, resulting in embryos heterozygous for *sna^MS2^*. For imaging of *Sna^Llama^*, *yw; P{w[+mC] = EGFP-STOP- bcd}* (hereafter named *bcd>*GFP^23^) was crossed to *His2A-RFPt/CyO*, followed by crossing of F1 virgin females to males bearing the *sna^Llama^* allele.

### Generation of CRISPR knock-ins and transgenic fly lines

Guide RNA sequences were selected using the CRISPR Optimal Target Finder site and cloned into pCFD3- dU6:3gRNA. The guide RNA sequences are listed in **Supplementary Table 3**. To create the *sna24xMS2* allele, a CRISPR recombination matrix comprised of a homology arm upstream of the 3’ UTR, a 24xMS2 stem loop sequence (derived from Bertrand et al., 1998), a floxed 3xP3-dsRed selection cassette and a downstream homology arm. The dsRed cassette was retained in *sna*^MS2^ stocks. To create the *sna^Llama^* allele, a CRISPR recombination matrix comprised of an 850bp genomic homology arm followed by the *sna* coding sequence, a flexible linker and *Drosophila*-optimized GFP-targeting nanobody, the genomic *Drosophila* 3’ UTR, a floxed 3xP3-dsRed selection cassette, and a 900bp genomic downstream homology arm. All genomic DNA was amplified using Phusion polymerase (Invitrogen), and the repair matrix was assembled in pBluescript-II SK(+). All matrices were sequenced prior to injection.

Generation of the *sna*^ΔATG^*/CyO-Hb>lacZ* line was accomplished using a ssODN co-CRISPR approach detailed in Levi et al^43^. Briefly, gRNAs (**Supplementary Table 3**) were constructed targeting the *sna* and *w* coding sequences via PCR with Phusion polymerase (Invitrogen) and assembled into pCFD4 *sna^ΔATG^_w^coffee^* using NEBuilderHiFi DNA Assembly Kit (New England Biolabs). A ssODN repair matrix targeting the *sna* N- terminal coding sequence was designed to add an EcoRI site in parallel for the screening. It was co-injected into *y^1^,M {vas-Cas9} ZH-2A* embryos along with a CRISPR repair matrix (pUC57-white [coffee] Addgene #84006) facilitating a conversion of the *w*^+^ allele into *w^coffee^* and pCFD4 *sna^ΔATG^_w^coffee^*. F0 flies were single- crossed to females bearing *sp/CyO-Hb>lacZ* and resulting F1 screened for the *w^coffee^* phenotype. F1 males were backcrossed to *sp/CyO-Hb>lacZ* balancer females and checked by genomic PCR and digestion (EcoRI) after several days to confirm the mutation. ssODNs were obtained from IDT Technologies.

The *sna^Distal^-24xMS2-y* and *sna^DistalCore^-24xMS2-y* minigenes have been previously described^44,45^. The *sna^DistalMut^* and *sna^DistalAlt^* enhancers were synthesized by Twist Biologicals. The *sna^DistalCore^* sequence was removed from pBPhi *snaShadowCore > snaPr> 24xMS2-y* using restriction enzyme-mediated excision and the *sna^DistalMut^* or *sna^DistalMut^* enhancer was inserted using NEBuilder HiFi DNA Assembly Kit (New England Biolabs) upstream of the *sna* promoter sequence. Enhancer sequences are listed in **Supplementary Table 4.** Transgenic flies were generated by PhiC31-mediated recombination into the VK33 locus (BL 9750). Injections were performed by the Drosophila Transgenesis Facility (Centro de Biología Molecular Severo Ochoa, Madrid) and FlyORF (Zurich). All stocks are homozygous with no observable viability defects, except for *sna*^ΔATG^*/CyO-Hb>lacZ* which is heterozygous. All lines are listed in **Supplementary Table 2**.

### Live Imaging

Embryos were permitted to lay for 2 h prior to mounting for live imaging. Embryos were hand- dechorionated using tape and mounted on a hydrophobic membrane prior to oil immersion to prevent desiccation, followed by the addition of a coverslip. Live imaging of MS2 embryos was performed with an LSM 880 with an Airyscan module (Zeiss). Z-stacks comprised of 30 planes with a spacing of 0.5 μm were acquired at a time resolution of 4.64s (*sna*^MS2^ and *sna* BAC reporters), 6.35s (*sog*^MS2^ and derivatives), or 3.86s (transgenic reporters) in fast Airyscan mode with laser power measured and maintained across embryos using a ThorLabs PM100 optical power meter (ThorLabs Inc.). All wild type background *sna*^MS2^ movies and *sna* BAC reporter movies were performed with the following settings: GFP excitation by a 488- nm laser (8uW with 10x objective) and RFP excitation by a 561 nm were captured on a GaAsP-PMT array with an Airyscan detector using a 40x Plan Apo oil lens (NA = 1.3) and a 2.5x zoom on the ventral region of the embryo centered (± 25 μm) on the presumptive ventral midline. Resolution was 640x640 pixels with bidirectional scanning. All *sog*^MS2^ (and derivative genotypes) movies were performed with the following settings: GFP excitation by a 488-nm (4.9uW with 10x objective) laser and RFP excitation by a 561 nm were captured on a GaAsP-PMT array with an Airyscan detector using a 40x Plan Apo oil lens (NA = 1.3) and a 2x zoom on the ventral/lateral region of the embryo including the ventral furrow. Time resolution was 6.35s and resolution was 800x800 pixels with bidirectional scanning. All RNAi- and overexpression-related *sna*^MS2^ movies were performed with the following settings: GFP excitation by a 488-nm laser (7.7uW with 10x objective) and RFP excitation by a 561 nm were captured on a GaAsP-PMT array with an Airyscan detector using a 40x Plan Apo oil lens (NA = 1.3) and a 2.5x zoom on the ventral region of the embryo centered (± 25 μm) on the presumptive ventral midline. Resolution was 640x640 pixels with bidirectional scanning. All *snail^Distal^* transgene (and derivative genotypes) movies were performed with the following settings: GFP excitation by a 488-nm laser (10.5uW with 10x objective) and RFP excitation by a 561 nm were captured on a GaAsP-PMT array with an Airyscan detector using a 40x Plan Apo oil lens (NA = 1.3) and a 3x zoom on the ventral region of the embryo centered (± 25 μm) on the presumptive ventral midline. For all imaging conditions, Airyscan processing was performed using 3D Zen Black v3.2 (Zeiss).

Imaging of *Sna^Llama^;His2A-RFP* was performed with an LSM880 (Zeiss). 18 Z-planes with a spacing of 1 μm were acquired with a time resolution of 42s/z stack. Movies were performed with the following settings: GFP excitation by a 488nm laser and RFP excitation by a 561nm laser captured on a GaAsP-PMT array using a 40x Plan Apo oil lens (NA = 1.3) at 1x zoom with resolution of 512x512 pixels. Laser power was measured and maintained across embryos using a ThorLabs PM100 optical power meter (ThorLabs Inc.).

### Image Analysis for MS2-MCP movies

The region of analysis was maintained at 25μm on either side of the presumptive ventral furrow. The intensity profile of the transcriptional sites imaged were extracted with a custom software developed in Python^TM^ ^46–48^ that has been previously published^16^ (<u>SegmentTrackv4.0</u>, https://github.com/ant-trullo/SegmentTrack_v4.0). However, for this study a post-processing tool was added (https://github.com/ant-trullo/SpotsFiltersTool). Transcription sites were infrequently resolved as two sister chromatids by the detection algorithm, potentially confounding distinguishing between sister chromatids and false detection events. A parameter was defined as the ratio between the convex hull surface determined by the two spots and their actual size. For sister chromatids this ratio is small (< 4) since the two spots are close, while a false detection event will generally be far from the real spot and with a small volume, so the ratio will be large (>4). Some blinking activation could potentially be falsely discarded with an overly stringent criteria for sequential frames showing activity, so a user-defined threshold was established for determining the number of sequential inactive time points required for detection to be considered false.

For analysis of homozygous *sna^MS2/MS2^* movies, a further post-processing tool was developed (https://github.com/ant-trullo/SistersSplitTool). To differentiate the alleles within the ‘spot’ signal, the position of each putative spot was identified relative to the centre of mass of the nucleus across time. These values were organized into two clusters using the Gaussian mixture algorithm from which the tracking in 3D was reconstructed. False detection events generally appeared far from the spot positions and as such could be removed as outliers based on their spatial position. Finally, a manual inspection and correction tool was implemented via a graphical user interface to perform detailed corrections.

### Single Molecule FISH and smFISH-immunofluorescence

Embryos heterozygous for the allele of interest were fixed in 10% formaldehyde/heptane for 25 minutes with shaking followed by storage in methanol at -20°C as previously described^21^. smFISH probes were designed and produced with primary labelling using Quasar 670 by LGC Biosearch Technologies Inc. Probe sequences are listed in **Supplementary Table 5.** smiFISH probes were designed following a modification of a previous methodology^49^ and produced by IDT.

Embryos were dehydrated with 2 × 5 min washes in 100% ethanol, followed by rehydration in PBT for 4 × 15 min and equilibration in 15% formamide/1 × SSC for 15 min. During equilibration, the probe mixture was prepared with a final concentration of 1 × SSC, 0.34 μg μL−1 E. coli tRNA (New England Biolabs), 15% formamide (Sigma), 5-μL probe, 0.2 μg μL−1 RNAse-free BSA, 2 mM vanadyl-ribonucleoside complex (New England Biolabs), and 10.6% dextran sulfate (Sigma) in RNAse-free water. The equilibration mixture was removed and replaced with probe mixture, and embryos were incubated overnight in the dark at 37°C with shaking. The following day, embryos were rinsed twice in equilibration mix and twice in PBT, followed by DAPI staining and three PBT washes before mounting in ProLong Gold mounting media (Life Technologies). For smFISH-IF, the same protocol was performed with addition of the primary antibody (rabbit anti-snail 1:500)^50^ in the probe mixture followed by secondary antibody (donkey anti-rabbit Alexa Fluor 488 1:500, Life Technologies) during PBT washes on the second day.

### Fixed Imaging

Fixed sample imaging was performed on an LSM 880 with an Airyscan module (Zeiss). Z-planes were acquired with 0.33 μm spacing to a typical depth of 25-30um from the apical surface of the embryo using laser scanning confocal in Airyscan super-resolution mode with a zoom of 3.0. DAPI excitation was performed with a 405nm laser, secondary Alexa Fluor 488 excitation with a 488nm laser (8μW), and Q670 with a 633nm laser (11.5μW), with detection on a GaAsP-PMT array coupled to an Airyscan detector. Airyscan processing was performed using 3D Zen Black v3.2 (Zeiss) prior to analysis. Embryos were staged based on membrane invagination.

### Single Molecule FISH Analysis

To analyse smFiSH data we used a custom software developed in Python^TM^ ^46–48^ that has been previously published^51^. Data was acquired in two channels, one for nuclei and the other for transcription, both in 3D (ZXY). Transcription channel was treated with a difference of Gaussian filter and the resulting image was thresholded. To find the optimal threshold value, the algorithm performed a systematic study over a range of different threshold values, followed by manual inspection and selection. The detected spots were composed of both transcriptional sites and single molecules that were further isolated into individual populations using a classifier and a visual tool for manual corrections when appropriate. The nuclei channel was pre-smoothed with a Gaussian filter and user-defined threshold individually for each z-slice to detect nuclei in 2D in each frame. These Z-frames were then combined in 3D to have a preliminary structure for nuclei. The following step was to find the smallest ellipsoid able to contain the detected 3D nucleus, which was then defined as the final nuclear volume. Finally, the intersection between the major axes of the ellipsoids was identified for each plane and used these points to simulate pseudo-cells with the Voronoi algorithm. Once the pseudo-cells were defined, the spatial position of transcription sites and single molecules was used to assign them to the appropriate ‘cell’. As each transcription site and single molecule has an associated intensity, the equivalent number of mRNA molecules was then calculated for each transcription site.

### Data Analysis of *snail*^Llama^

Visualization and analysis of the time series data was performed using custom software developed in Python™ ^46–48^ enabled by a graphical user interface (NucleiTracker3D, https://github.com/ant-trullo/NucleiTracker3D). Raw data consisting of a two channel TZXY series with Sna^Llama^-GFP intensity and His2A-mRFP as a reference nuclear marker. The reference nuclear channel was pre-smoothed using a Gaussian filter, and then thresholded with an Otsu algorithm. The resulting connected components were labelled in 3D treated with a 3D watershed algorithm to separate touching nuclei. Hyper-segmented nuclei were recognized using a classifier algorithm previously trained to identify ‘sub-nuclear’ fragments and combine them with neighbouring nuclei, privileging combinations with the fewest sub-components. Segmented nuclei were tracked by sequentially projecting the *t-1* time point nuclear mask onto the frame of time point *t* and tagging each nuclei *n* with the most coincident nuclear tag of the projected mask using a median filter, as nuclear motion between timepoints *t-1* and *t* was less than the nuclear radius in XYZ. This 3D-tracked nuclear mask was then projected on the Sna^Llama^-GFP channel to retrieve the nuclear GFP fluorescence. For each time point the average nuclear out-of-pattern and in-pattern GFP signal intensity was retrieved. An enrichment ratio was then calculated by dividing the in-pattern by the out-of-pattern GFP signal intensity in a time-dependent manner^52^.

### Deconvolution analysis of the data using BurstDeconv

We use BurstDeconv^14^ to obtain, for each transcription site, the sequence of processive transcription initiation events. The method considers that the single site MS2 signal is a sum of identical single polymerase contributions translated in different positions corresponding to the initiation events. The signal is calibrated in terms of numbers of polymerases using smFISH^16^, hence the amplitude of the signal polymerase contribution is equal to one. The dwell time, defined as the duration of the single polymerase contribution signal, is obtained using the autocorrelation function of the stationary single site MS2 signal in nc13^14,17^. Finally, the processive transcription initiation events positions are obtained by combining a genetic algorithm with a local optimisation procedure to minimise the least squares distance between the predicted and observed signal. One should note that this procedure, initially applied to stationary signals^14–16^, can also be applied without modification to non-stationary signals.

### Multi-exponential regression and multiple state transcription model reverse engineering using the survival function on stationary segments of the signal

The single-site MS2 signal can be approximated as stationary within segments corresponding to two regimes: the fully unrepressed regime during nuclear cycle 13 and the fully repressed regime during the terminal segment of the nuclear cycle 14 signal. On such segments we use the Meyer-Kaplan method to compute a survival function, representing the nonparametric cumulative distribution function of the waiting times separating successive transcription events. A multi-exponential fitting of the survival function using N exponentials (N=2,3 for two and three states models, respectively) is followed by the reverse engineering of the transcription model using a symbolic method described previously^14,19^. The multi-exponential regression determines the number of states based on parsimony. We select the simplest model that provides a good fit to the survival function, as judged by three criteria: the least-squares objective function, the confidence interval of the Kaplan-Meier estimate, and the Kolmogorov-Smirnov test.

### Determining the dwell time from autocorrelation

The signal autocorrelation function is defined as 𝑅(𝑡, 𝑡^′^) = 𝐶𝑜𝑣(𝑥(𝑡), 𝑥(𝑡^′^)), where 𝑥(𝑡) is the single site MS2 signal. For a stationary MS2 signal, this function depends only on 𝜏 = 𝑡 − 𝑡^′^ according to the relation:

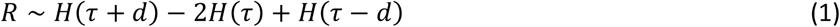

where 𝑑 is the dwell time, 𝐻(𝑥) = −𝑥𝜃(−𝑥) and

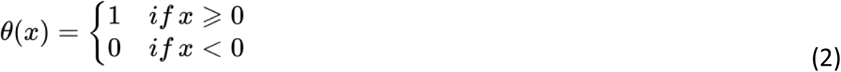

is the Heaviside function. A derivation of (1) has been previously published^17^.

For the purposes of this research, the stationary signal from nuclear cycle 13 was used to fit the dwell time (**Supplementary Figure 8**).

### Estimating the time-dependent repression

The MS2 signal is non-stationary during the nuclear cycle 14. Typically, we notice a decrease of the signal amplitude, suggesting increasing repression. To characterize the time dependence of the repression, we use a moving window method to estimate the time dependence of the mean waiting time *<τ>* between successive processive transcription initiation events. As shown elsewhere^19^, there is a general formula relating the mean *<τ>* and *p_ON_ k_ini_*, where *p_ON_* is the probability of the transcribing (ON) state and *k_ini_* is the transcription initiation rate in this state. This formula is valid for all finite state Markov models, irrespective of their number of states. We reproduce here the reasoning leading to this formula. The mean number of transcription events on an interval *[0,T]* is *T/<τ>*. The same number is equal to *T·p_ON_·k_ini_*, because in the state ON, the promoter initiates with intensity *k_ini_* and the total time spent in the ON state is *T· p_ON_*.

It follows that:

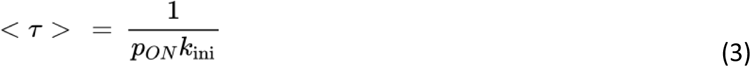

For a stationary signal *p_ON,_ k_ini_* and thus *<τ>* are functions of time. For increasing repression, *<τ>* is increasing (the initiation events are rarer). Therefore, the estimate of *<τ>* and implicitly that of *p_ON_k_ini_* is a method to gauge repression. To estimate *<τ>*, we define a narrow moving window centered on successive time frames and consider all the waiting times from all transcription sites contributing to signal observed in the moving window, gathering sites observed in several movies for enriched statistics. The width of the windows is 5-8 frames, i.e. 22.7-36.3 seconds for *sna*. This width is enough for including a sufficiently large number of waiting times for an accurate estimate of the mean *<τ>*. We compute the uncertainty bounds of the mean for each movie independently using the central limit theorem with a 95% confidence interval, and then plot the minimum lower bound and the maximum upper bound over all the movies.

### Bayesian Change Point Detection for determining the onset of repression

The BCPD method^22^ is used to determine the onset of repression by determining the probability of having a change point denoting a sudden change in the parameters that generate the data. The distribution of the run length 𝑟(𝑡), defined as the time since the most recent change point, is learned from the data using this method. At each time step, 𝑟(𝑡) increases by 1 if there is no change in the distribution, or it returns to zero when there is a change with a certain probability. It is based on a recursive message-passing algorithm for the joint distribution of observations and run lengths. The algorithm assumes that: i) the single nucleus MS2 signal follows a normal distribution with unknown mean and variance, and ii) the run length advances without memory, according to a geometric distribution.

An illustration of the BCPD method is given in **Supplementary Figure 4**. For discretized times 𝑡 = 1,2, … the run length 𝑟(𝑡) is defined by the following relation:

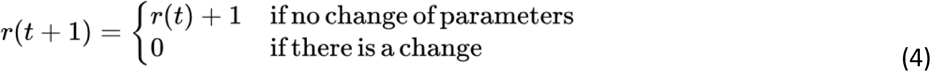

The BCPD method computes the conditional probability of the run length, given the observed values of the signal. The prior of this distribution is learned from the data. We have tested the method using synthetic data generated using the Gillespie algorithm and a two-state telegraph model. The model includes an RNA and a protein pool and considers autorepression by considering that the kinetic parameters depend on the protein level according to decreasing and increasing Hill functions, respectively (**Supplementary Figure 4A**). We have used the promoter transcription initiation events to compute a synthetic MS2 signal for each site. The traces generated by this model show change points corresponding to onset of the repression. These change points are correctly detected by the method as shown visually in **Supplementary Figure 4A** where we confirm that the switching parameters are stable after the change point is found. Since this is a Bayesian approach, the change point is identified with a probability. A threshold was imposed such that the probability p must be > 0.8 for the change to be retained (**Supplementary Figure 4B** middle vs bottom panel). Three further criteria for detecting the change point were applied to change point detection on experimental data. First, due to the noisy nature of the data, a smoothing criterion was applied. The smoothing process involves convoluting the data with a box filter kernel of size 2 so that the rapid fluctuations in the experimental data are attenuated in the smoothed data. Second, a changepoint detection window was imposed such that the selected changepoint was the first change point to occur after 60% of the maximum to eliminate changepoints during the post-mitotic activation period. Third, a minimal filtering criterion was applied to individual traces. The filtering process includes the elimination of signals in the attenuated part that are either 0 everywhere or have less than 5% of the mean number of initiation events before the identified changepoint.

The part of the MS2 signal after the repressor onset checkpoint in nc14 and the full MS2 signal in nc13 were then used for inferring promoter models for the repressed and active phases, respectively, using BurstDeconv^14^. Model selection was performed using three fitting scores: the objective function, the confidence interval of the empirical survival function and the Kolmogorov-Smirnov test comparing the empirical and predicted distribution of waiting time between successive transcription events^14^.

### Homozygous *sna^MS2/MS2^* data analysis

For *sna^MS2/MS2^* movies, deconvolution of the data was processed in two separate approaches: 1) the alleles were segregated based on the first (1^st^ allele - paired) and second (2^nd^ allele - paired) activated allele in a single nucleus, or 2) all alleles were pooled regardless of their ‘mother’ nucleus or activation order and randomly allocated to one of two pools (“1^st^ allele - random” or “2^nd^ allele - random”). For the paired allele pools, alleles were retained only if both alleles in a nucleus entered stable repression to avoid bias, which was >95% of nuclei analyzed. For the randomly assigned pools, all alleles that entered stable repression were retained.

### The Hill equation model for the mean mRNA production rate

The key to this model is the nonlinear relation between the repressor protein concentration and the mRNA production rate:

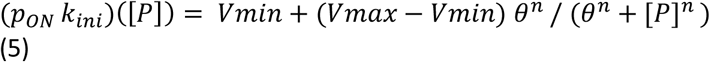

where [P] is the repressor protein concentration, *Vmax* and *Vmin* are the maximal and minimal mRNA production rates, 𝜃 is the threshold repressor protein concentration at half-maximal repression, and *n* is the Hill coefficient. As suggested in its first utilisation by Hill^25^, *n* does not necessarily take integer values. Values *n* > 1 indicate positive cooperativity, meaning that binding of one molecule of repressor to DNA facilitates the binding of additional molecules and/or enhances their repressive effect.

If the protein level and the product 𝑝_𝑂𝑁_ 𝑘_𝑖𝑛𝑖_ (also described in text as the inverse of *<τ>*, see Eq.(3)) are both known, Eq.(5) can be used to estimate the parameters *V*, *θ*, and *n*, with no assumption on the identity of the repressor or mechanism of repression.

### Stochastic model of transcriptional repression

The Hill equation model for the mean mRNA production is useful for a first analysis of the data. It has the advantage of estimating a Hill coefficient which is a measure of cooperativity. This model does not propose mechanisms or dynamics for transcriptional repression. Therefore, we need a dynamical model of the transcriptional process under repression. Furthermore, the Hill equation cannot explain the stochastic fluctuations in mRNA production. Using the BCPD method, we extracted repression onset times from experimental data and observed that these times vary across different nuclei. A stochastic model is therefore needed to gain more insight into the distribution of these times.

We therefore propose a discrete state transcription model for the endogenous promoter, in which the transcription dynamics is described by the stochastic transition between states. To account for the effect of the repressor on transcription in the simplest way, we consider that the transition rates between discrete promoter states are modulated by the repressor in a Hill-dependent manner.

The objectives of this mathematical model are to 1) check whether our model can reproduce the distribution of repression onset times extracted using BCPD from the experimental data in nc14 and 2) predict the effect of cooperativity on coordinated repression (by scanning over different ranges of Hill coefficient and threshold values).

To build this model, we consider the following experimental observations:

1. The BurstDeconv analysis of stationary segments of the single nucleus MS2 signal indicates a two-state model for nc13 and a three-state model for the last segment of nc14 during repression. The three-state model includes two OFF states: a long one, denoted OFF1, and a short one, denoted OFF2. This suggests a transition from a two-state system to a three-state system during nc14. We do not know *a priori* which of the states (OFF1 or OFF2) is newly introduced and which is a continuation of the OFF state from nc13/early nc14.
2. The transgene experiments show that when the number of Sna binding sites and therefore the repression is weakened, there is a transition from a three state model to a two state model. This is consistent with observation 1.
3. The BurstDeconv analysis of different genotypes suggest that the switching rate constants between discrete states in the three state model depend on repression as follows (see **Supplementary Figure 10A-G**):

a. k_1_^p^ = 1/T_OFF1_ decreases strongly with repression,
b. k_1_ increases strongly with repression, with k _m_ being very small (its inverse corresponding to hours) when repression is very weak,
c. k_2_ increases strongly with repression,
d. k_2_ = 1/T_OFF2_ has a mild increase with repression,
4. k_ini_ is very similar for nc13 and the last segment of nc14.

According to 3b, the long state OFF1 is very rare (practically not accessible) at weak repression, and therefore the long OFF1 likely corresponds to the new state induced by repression.

Here we hypothesized that weakening the repression by eliminating Sna binding sites in transgenes is equivalent to decreasing the concentration of Sna for endogenous promoters. We do not exclude the possibility that other TFs (activators or repressors) may also modulate the expression from the endogenous locus, but we consider that the main contribution to the observed variation of parameters is due to changes in the Sna concentrations.

Thus, the transcriptional process is described as a three state Markovian model and governed by the following set of chemical reactions:

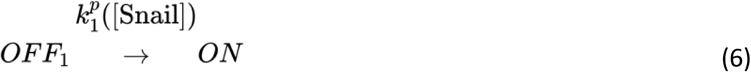

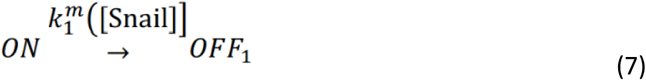

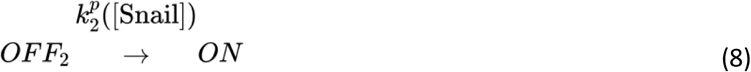

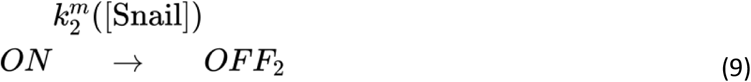

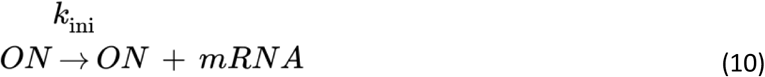

The Hill-like dependence of parameters 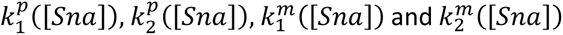 is given by the equations (we use increasing and decreasing Hill functions for the ON-> OFF, and OFF-> ON transitions, respectively):

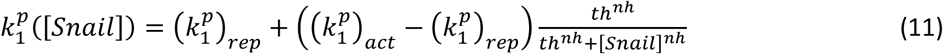

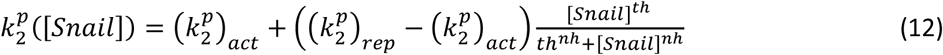

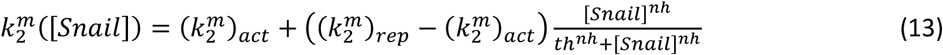

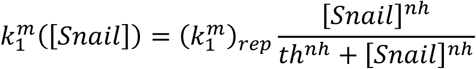

where 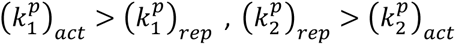 are OFF-> ON rates, 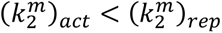 are ON->OFF rates in active and repressed phases, respectively.

The rate parameters 𝑘_𝑖𝑛𝑖_,(𝑘_2_^𝑚^ )_𝑎𝑐𝑡_,(𝑘_2_^𝑝^ )_𝑎𝑐𝑡_,(𝑘_2_^𝑚^ )_𝑟𝑒𝑝_,(𝑘_2_^𝑝^ )_𝑟𝑒𝑝_,(𝑘_1_^𝑚^ )_𝑟𝑒𝑝_,(𝑘_1_^𝑝^ )_𝑟𝑒𝑝_ were chosen equal to the already estimated values, in the active nc13 and repressed nc14 regimes (**Supplementary Table 1**, 2 state and 3 state non-sequential model). All parameters except 𝑘_𝑖𝑛𝑖_ depend on [𝑆𝑛𝑎].

The model remains with three free parameters *nh*, *th*,(𝑘_1_^𝑝^ )_𝑎𝑐𝑡_ that were fitted to the *p_ON_·k_ini_* vs [Sna] curves and to the distribution of repression onset times data. The result of the fit is given in the table below:

| Symbol | Value | Justification |
| --- | --- | --- |
| $k_{ini}$ | 0.35 s<sup>-1</sup> | nc13 and nc14 last segment value |
| $(k_2^m)_{act}$ | 0.007 s<sup>-1</sup> | nc13 value |
| $(k_2^p)_{act}$ | 0.011 s<sup>-1</sup> | nc13 value |
| $(k_1^p)_{rep}$ | 0.001 s<sup>-1</sup> | nc14 last segment value |
| $(k_1^m)_{rep}$ | 0.0075 s<sup>-1</sup> | nc14 last segment value |
| $(k_2^m)_{rep}$ | 0.11 s<sup>-1</sup> | nc14 last segment value |
| $(k_2^p)_{rep}$ | 0.045 s<sup>-1</sup> | nc14 last segment value |
| $(k_1^p)_{act}$ | value s<sup>-1</sup> | free parameter, fitted |
| nh | value | free parameter, fitted |
| th | value | free parameter, fitted |

The Gillespie algorithm was used to numerically simulate the model. To account for post-mitotic lag in transcriptional reactivation, we assumed that transcription began after a lag time in nc14, which varied for each nucleus. The simulated model distribution of lag times was sampled from the experimental post-mitotic lag duration of the *sna* gene, which was obtained using the BurstDeconv deconvolution output. In these simulations, the protein signal was considered deterministic; in other words, different nuclei receive regulatory cues that depend on the time point only.

A synthetic MS2 signal was computed for each simulation using the dwell time parameter of the *sna*^MS2^ gene such that the simulated data can be compared with the experimental results in nc14.

In order to test the dependence of the repression onset time on the Hill function parameters, 154 value pairs of {*(k_1_ )_act_*, *nh, th*} were benchmarked with each simulation run for 484 sample traces and a duration of 30.93 min to match the time window studied for the expression of the experimental *sna*^MS2^ gene. Each simulation was then run for each set of {*(k_1_ )_act_, nh, th*} through the BCPD algorithm to obtain the repression onset time. No smoothing was applied to the signal used in the stochastic model since no noise was added. **Figure 6C** shows the results of the best fit simulations. We then chose the best values of {*(k_1_ )_act_*, *nh, th*} that fit the data according to the sum of squared error between *p_ON_·k_ini_* of the data and the simulations. **Figure 6D** shows the distribution of the repression onset time time computed for these parameters. The good agreement between predicted and experimental distributions validates the stochastic model.

### qPCR analysis

To test changes in expression of pause-related genes, 0–2 h embryos were homogenized in Trizol (Invitrogen), and RNA was extracted as directed by the manufacturer. Reverse transcription was performed using Superscript IV (Invitrogen) with random hexamers. Measurements were performed in biological and technical triplicate. qPCR analysis was performed using LightCycler 480 SYBR Green I Master Mix (Roche) using primers listed in **Supplementary Table 6**. Analysis was performed using Microsoft Excel and Prism (Graphpad 9.1.1).

### Snail binding site identification

Sna ChIP data was obtained from previously published data (GSE68983)^24^. To identify potential transcription factor binding sites, we employed the FIMO (Find Individual Motif Occurrences) tool^53^ with Sna motifs obtained from the JASPAR database^54^. The significance threshold for motif matches was set at p<1e-3. Sna binding sites were queried in the local neighbourhood of the highest (>500) ChIP signal for Sna.

---

**Supplementary Figure 1:** Profiling *sna^MS2^*. A) *sna^MS2/+^* heterozygotes showing coincident labelling of *sna^MS2^* allele and endogenous *snail* mRNA at the transcription site (inset). Scale bar represents 10 mm. B) Sample membrane invagination (arrowheads) indicating partitioning of embryos into early (left) and late (right) nc14. C) Activation profiles for individual *sna^MS2/+^* movies during nc14. D) Average intensity of actively transcribing nuclei for individual *sna^MS2/+^* movies. Statistics: *sna^MS2/+^* nuclear cycle 14: N=6 embryos, n=484 nuclei.

**Supplementary Figure 2:** Generation of *sna^ΔATG^/CyO-Hb>lacZ* line. A) schematic of co-CRISPR strategy targeting *sna* and *w* simultaneously. B) Crossing scheme to recover mutant *sna^ΔATG^* alleles. Adapted from Levi *et al*.^56^ C) Sequencing results of fly. Arrows indicate mutations relative to wild type sequence. D) smFISH-IF to show co-occurrence of *sna* transcription with concomitant expression of Sna protein in *yw* control and absence of Sna protein in *sna^ΔATG/ΔATG^*.

**Supplementary Figure 3:** Autocorrelation of the *sna* transcription signal in nuclear cycle 13. A) Sample trace demonstrating autocorrelation of a *sna* transcription trace in nc14. B) Dwell time of the *sna^MS2^* signal in nuclear cycles 13 and 14 compared to the theoretical prediction for a polymerase speed of 25bp/s and a retention time at the transcription site of 0s.

**Supplementary Figure 4:** Bayesian Change Point Detection (BCPD) identifies active and repressed periods. A) Synthetic data generated using the Gillespie algorithm with a two-state auto-repressive model. In this model, population-level 𝑘_𝑂𝑁_ and 𝑘_𝑂𝐹𝐹_ depend on the protein level according to decreasing and increasing Hill functions, respectively (top). Synthetic traces were computed with a detected change point highlighted by a change in colour (middle) and a global change in kinetic parameters was observed (bottom). B) Synthetic traces generated using the Gillespie algorithm with their respective run length identified by the BCPD algorithm and classification into active and repressed states. C) Sample synthetic trace with no change in activation/repression is correctly detected by the BCPD algorithm. D) Cumulative distribution of BCPD algorithm-identified changepoints for synthetic data compared to the repression identified by the Hill algorithm.

**Supplementary Figure 5:** Dual allele labelling of *sna^MS2/MS2^* shows no difference in repression onset between alleles. A) Schematic of ‘paired’ allele assignment dividing alleles in the same nucleus into separate analysis pools based on initial activation order. B) Schematic of ‘random’ allele assignment where alleles were randomly sorted into two equal sized pools irrespective of nucleus or activation order. C-D) Distribution of repression onset for first active allele (C) or second active allele (D). E-F) Distribution of repression onset for each of the randomly assigned pools. Statistics: *sna^MS2MS2+^* paired alleles: N=3 embryos, n=228 nuclei per class; *sna^MS2MS2+^* randomly assigned alleles: N=3 embryos, n=255 nuclei (first allele) and n=256 nuclei (second allele).

**Supplementary Figure 6:** Survival functions for *sna^Distal^* transgene series in nc14. A-B) Survival function of the distribution of waiting times between polymerase initiation events (red circles) for *snail^Distal^* with the two-exponential fitting (A) or three-exponential fitting (B) of the population estimated using the Kaplan– Meyer method (black line). The dashed lines indicate 95% confidence interval. A red cross indicates a rejected fitting. A green check indicates an accepted fitting. C-D) Survival function of the distribution of waiting times between polymerase initiation events (red circles) for *snail^DistalAlt^* with the two-exponential fitting (C) or three-exponential fitting (D) of the population estimated using the Kaplan–Meyer method (black line). The dashed lines indicate 95% confidence interval. A red cross indicates a rejected fitting. A green check indicates an accepted fitting. E) Survival function of the distribution of waiting times between polymerase initiation events (red circles) for *snail^DistalMut^* showing a two-exponential fitting of the population estimated using the Kaplan–Meyer method (black line). The dashed lines indicate 95% confidence interval. A green check indicates an accepted fitting. F) Survival function of the distribution of waiting times between polymerase initiation events (red circles) for *sna^DistalCore^* showing a two-exponential fitting of the population estimated using the Kaplan–Meyer method (black line). The dashed lines indicate 95% confidence interval. A green check indicates an accepted fitting. Statistics: *snail^Distal^* N=3 embryos, n=224 nuclei; *snail^DistalAlt^* N=5 embryos, n=220 embryos; *snail^DistalMut^* N=2 embryos, n=145 nuclei; *snail^DistalCore^* N=3 embryos, N=194 nuclei.

**Supplementary Figure 7:** *sna* expression in nc14 is stationary only during stable repression. A-B) Kinetic parameter stability as a function of time in nc14 during the first 30 minutes (A) or during stable repression only (B). Transcription expressed as the product of the probability to be active (*p_ON_*) and the RNA polymerase II initiation rate (*k_ini_*). C-D) Survival function of the distribution of waiting times between polymerase initiation events (red circles) for stable repression of *sna* with the two-exponential fitting (C) or three-exponential fitting (D) of the population estimated using the Kaplan–Meyer method (black line). The dashed lines indicate 95% confidence interval. A red cross indicates a rejected fitting. A green check indicates an accepted fitting. Statistics: *sna^MS2/+^* nuclear cycle 14: N=6 embryos, n=484 nuclei.

**Supplementary Figure 8:** *sna* expression in nc13 is in a fully active two-state regime. A) Instantaneous activation percentage (mean ± SEM) curves of ventral nuclei during nc13. Time zero is from anaphase during nc12-nc13 mitosis. B) Fluorescence intensity of actively transcribing nuclei (mean ± SEM) during nc13. Time zero is from anaphase during nc12-nc13 mitosis. C) Kinetic parameter stability as a function of time in nc13. Transcription expressed as the product of the probability to be active (*p_ON_*) and the RNA polymerase II initiation rate (*k_ini_*). D) Heatmap showing the number of polymerase initiation events in nc14 for *sna* in nc13 as a function of time. Each row represents one nucleus, and the number of Pol II initiation events per 30 s bin is indicated by the bin color. E) Survival function of the distribution of waiting times between polymerase initiation events (red circles) for *sna* showing a two-exponential fitting of the population estimated using the Kaplan–Meyer method (black line). The dashed lines indicate 95% confidence interval. A green check indicates an accepted fitting. F) Representation of estimated bursting dynamics for *sna* in nc13. Permissive ON state durations are depicted in green and inactive OFF states in red and orange, and probabilities of each state shown above. Statistics: *sna^MS2/+^* nuclear cycle 13: N=4 embryos, n=274 nuclei.

**Supplementary Figure 9:** Pausing is not rate-limiting during *sna* stable repression. A) schematic describing different pausing-related controls mediated by the Paf1 complex and pTEF-B (Cyclin T/CDK9) complex. B-C) qPCR quantification of *paf1* knockdown (B) or *Cyclin T* over-expression (C) relative to indicated controls. D-K) Survival function of the distribution of waiting times between polymerase initiation events (red circles) for *white-RNAi* (D,E), *paf1-RNAi^A^* (F,G) *paf1-RNAi^B^* (H,I), and *Nelf-A RNAi* (J,K) with a two- (upper row) or three-exponential (lower row) fitting of the population estimated using the Kaplan–Meyer method (black line). L-O) Survival function of the distribution of waiting times between polymerase initiation events (red circles) for *Gal4>+* (L,M) and *UAS:CycT* (N,O) with a two- (upper row) or three-exponential (lower row) fitting of the population estimated using the Kaplan–Meyer method (black line). For all fittings, the dashed lines indicate 95% confidence interval. A green check indicates an accepted fitting. Statistics: *white-RNAi* > *sna^MS2/+^*: N=2 embryos, n=183; *paf1-RNAi^A^* > *sna^MS2/+^*: N=2 embryos, n=177; *paf1- RNAi^B^*> *sna^MS2/+^*: N=2 embryos, n=184; *Nelf-A RNAi*> *sna^MS2/+^* N=2 embryos, n=154; *Gal4>+*> *sna^MS2/+^*: N=2 embryos, n=191 nuclei; *UAS:CycT> sna^MS2/+^*: N=2 embryos, n=154 nuclei.

**Supplementary Figure 10:** Comparison of kinetic parameters for endogenous *sna* and *sna^Distal^* transgene mutant series. A-B) Schema of two state (A) and three state (B) topology with transitions indicated. For two-state genotypes, the k^1^_m_ and k^1^_p_ transitions cannot be reached. C-G) Comparison of state transition rates for k^1^_m_ (C), k^1^_p_ (D), k^2^_m_ (E), k^2^_p_ (F), and the RNA production rate (G). Genotypes are arranged in order of decreasing repression.

**Supplementary Table 1:** Kinetic parameters for promoters derived from deconvolution and multi-exponential regression fitting of live imaging data. Minimum and maximum values indicate the boundaries of the error interval. State durations are calculated from the provided switching rates (*k_i_* ) and time durations for each state are provided as ‘*T(state)*’. State probability values are indicated as ‘*p(state)*’. Bold indicates the most parsimonious appropriate fitting of the data. The table also provides the objective functions and one-sided Kolmogorov-Smirnov test results.

**Supplementary Table 2:** *Drosophila* lines used in this manuscript.

**Supplementary Table 3:** guide RNA and ssODN sequences used to generate *sna*^MS2^, *sna^ΔATG^/CyO-Hb>lacZ*, and *Sna^Llama^* CRISPR alleles.

**Supplementary Table 4:** Enhancer sequences for *snail^Distal^* transgenes, related to Figure 5.

**Supplementary Table 5:** Single molecule fluorescent *in situ* hybridization probes for endogenous *sna* (related to Figure 1).

**Supplementary Table 6:** qPCR primers related to Supplementary Figure 9.

**Supplementary Movie 1:** Description: Live imaging of *sna*^MS2^ representative of nc13-14 beginning at mitosis. Nuclei are detected using His2Av-mRFP and MS2 using MCP-GFP.

**Supplementary Movie 2:** Description: Live imaging of *sog*^MS2^ representative of NC14 beginning at mitosis. Nuclei are detected using His2Av-mRFP and MS2 using MCP-GFP.

**Supplementary Movie 3:** Description: Live imaging of *sna*^Llama^ representative of nc13 and nc14 beginning at mitosis. Nuclei are detected using His2Av-mRFP.

**Supplementary Movie 4:** Description: live imaging of *sna^Distal^-24xMS2-y* representative of nc14 beginning at mitosis. Nuclei are detected using His2Av-mRFP and MS2 using MCP-GFP.

**Supplementary Movie 5:** Description: live imaging of *sna^DistalAlt^-24xMS2-y* representative of nc14 beginning at mitosis. Nuclei are detected using His2Av-mRFP and MS2 using MCP-GFP.

**Supplementary Movie 6:** Description: live imaging of *sna^DistalMut^-24xMS2-y* representative of nc14 beginning at mitosis. Nuclei are detected using His2Av-mRFP and MS2 using MCP-GFP.

**Supplementary Movie 7:** Description: live imaging of *sna^DistalCore^-24xMS2-y* representative of nc14 beginning at mitosis. Nuclei are detected using His2Av-mRFP and MS2 using MCP-GFP.

