## Supplementary Figures 1-10 and Tables 1-6 for "Coordinated active repression operates via transcription factor cooperativity and multiple inactive promoter states in a developing organism"

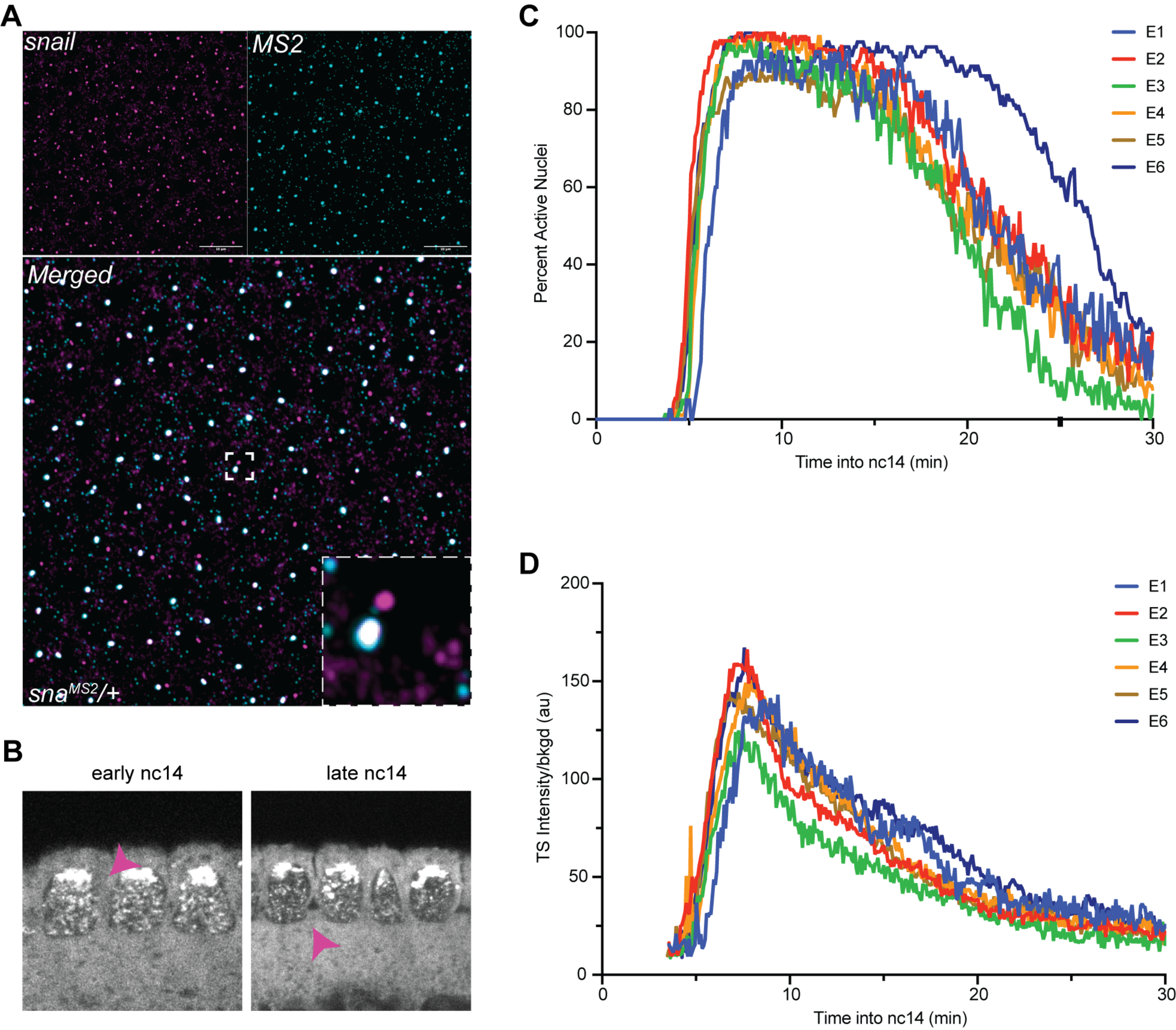
A
snail
MS2
Merged
sna^MS2/+
B
early nc14
late nc14
C
Percent Active Nuclei
Time into nc14 (min)
E1
E2
E3
E4
E5
E6
D
TS Intensity/bkgd (au)
Time into nc14 (min)
E1
E2
E3
E4
E5
E6

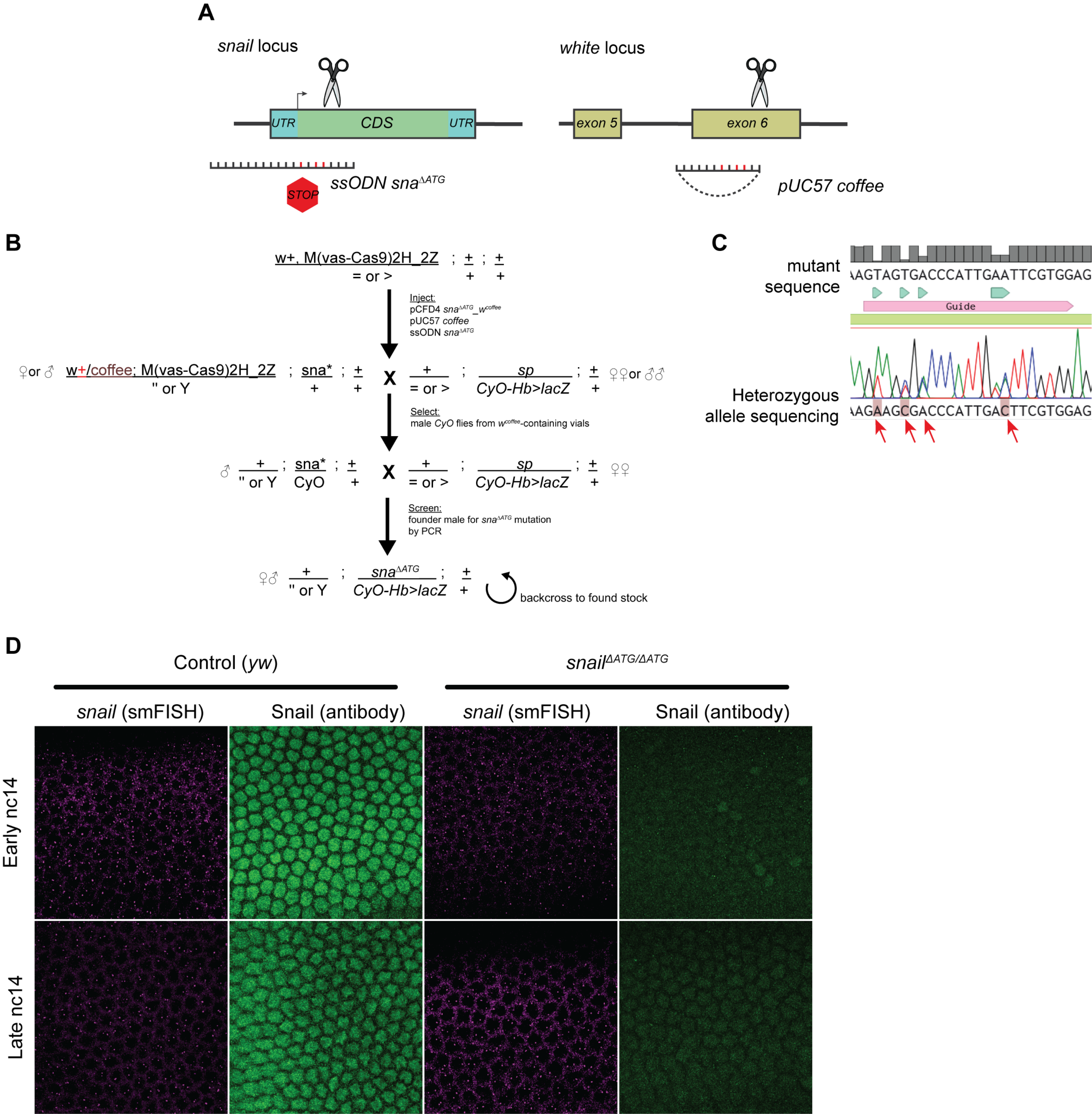
A
snail locus
UTR
CDS
UTR
ssODN sna^ΔATG
STOP
white locus
exon 5
exon 6
pUC57 coffee
B
w+, M(vas-Cas9)2H_2Z ; + ; +
= or > ; + ; +
Inject:
pCFD4 sna^ΔATG_w^coffee
pUC57 coffee
ssODN sna^ΔATG
♀or ♂ w+/coffee; M(vas-Cas9)2H_2Z ; sna* ; +
" or Y ; + ; +
X
+ ; sp ; +
= or > ; CyO-Hb>lacZ ; +
♀♀or ♂♂
Select:
male CyO flies from w^coffee-containing vials
♂ + ; sna* ; +
" or Y ; CyO ; +
X
+ ; sp ; +
= or > ; CyO-Hb>lacZ ; +
♀♀
Screen:
founder male for sna^ΔATG mutation
by PCR
♀♂ + ; sna^ΔATG ; +
" or Y ; CyO-Hb>lacZ ; +
backcross to found stock
C
mutant
sequence
AAGTAGTGACCCATTGAATTCGTGGAG
Guide
Heterozygous
allele sequencing
AAGAAGCGACCCATTGACTTCGTGGAG
D
Control (yw)
snail^ΔATG/ΔATG
snail (smFISH)
Snail (antibody)
snail (smFISH)
Snail (antibody)
Early nc14
Late nc14

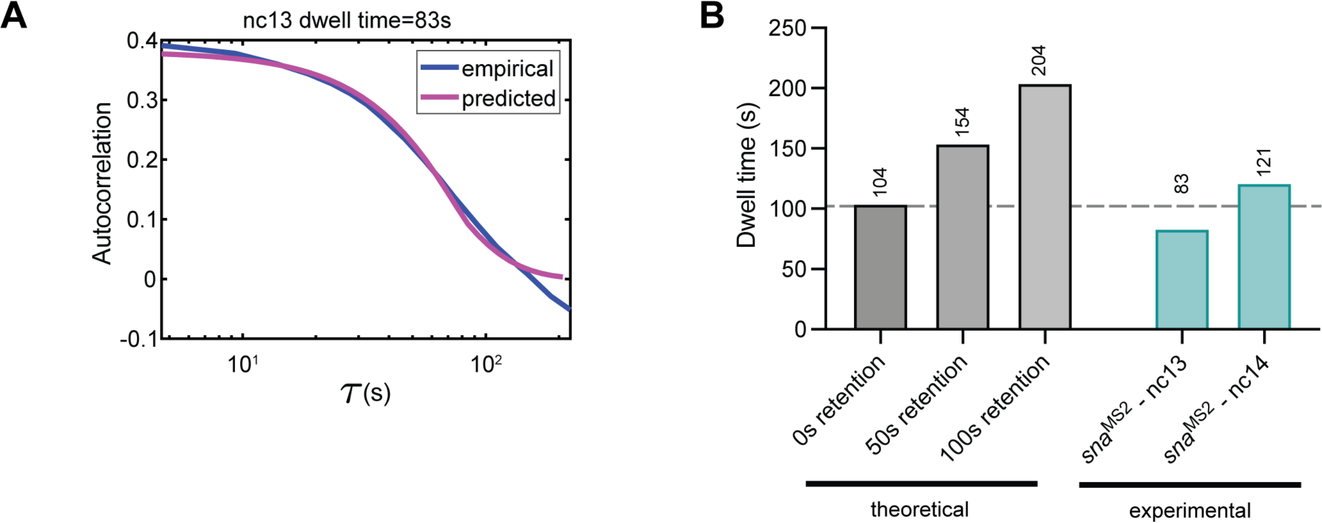
A
nc13 dwell time=83s
Autocorrelation
0.4
0.3
0.2
0.1
0
-0.1
empirical
predicted
$10^1$
$10^2$
$\tau$ (s)
B
Dwell time (s)
250
200
150
100
50
0
104
154
204
83
121
0s retention
50s retention
100s retention
$sna^{MS2}$ - nc13
$sna^{MS2}$ - nc14
theoretical
experimental

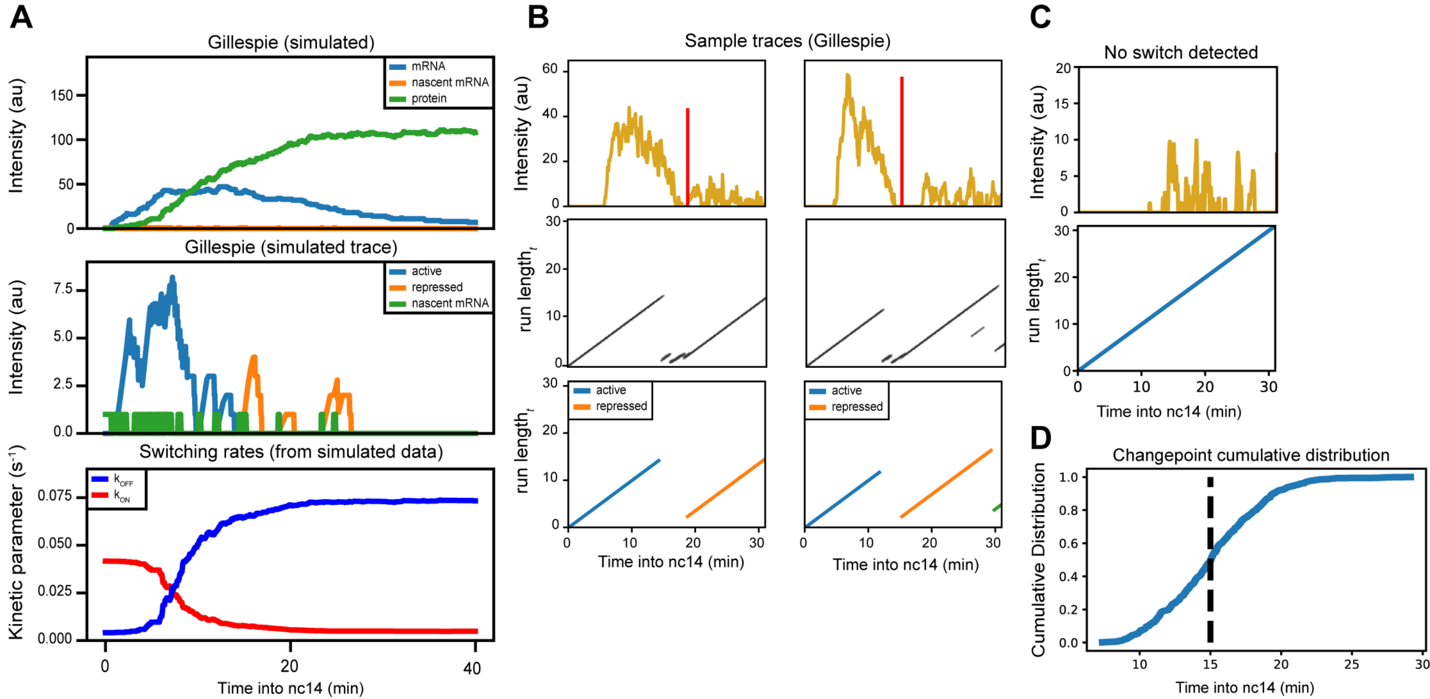

A
Gillespie (simulated)
mRNA
nascent mRNA
protein
Intensity (au)
Gillespie (simulated trace)
active
repressed
nascent mRNA
Intensity (au)
Switching rates (from simulated data)
k_OFF
k_ON
Kinetic parameter (s^-1)
Time into nc14 (min)
B
Sample traces (Gillespie)
Intensity (au)
run length_t
active
repressed
Time into nc14 (min)
C
No switch detected
Intensity (au)
run length_t
Time into nc14 (min)
D
Changepoint cumulative distribution
Cumulative Distribution
Time into nc14 (min)

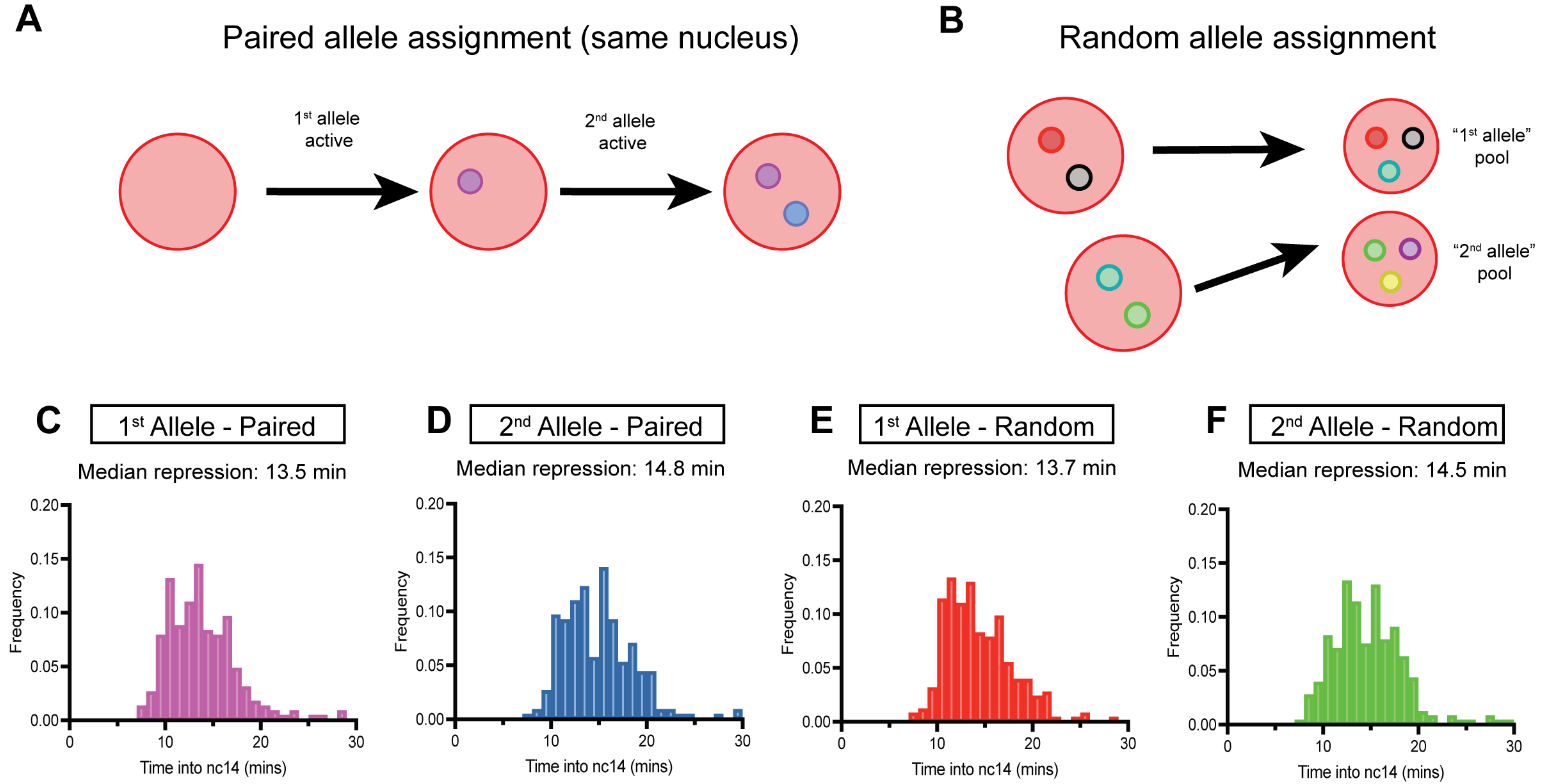
A
Paired allele assignment (same nucleus)
1st allele active
2nd allele active
B
Random allele assignment
"1st allele" pool
"2nd allele" pool
C
1st Allele - Paired
Median repression: 13.5 min
D
2nd Allele - Paired
Median repression: 14.8 min
E
1st Allele - Random
Median repression: 13.7 min
F
2nd Allele - Random
Median repression: 14.5 min
Frequency
Time into nc14 (mins)

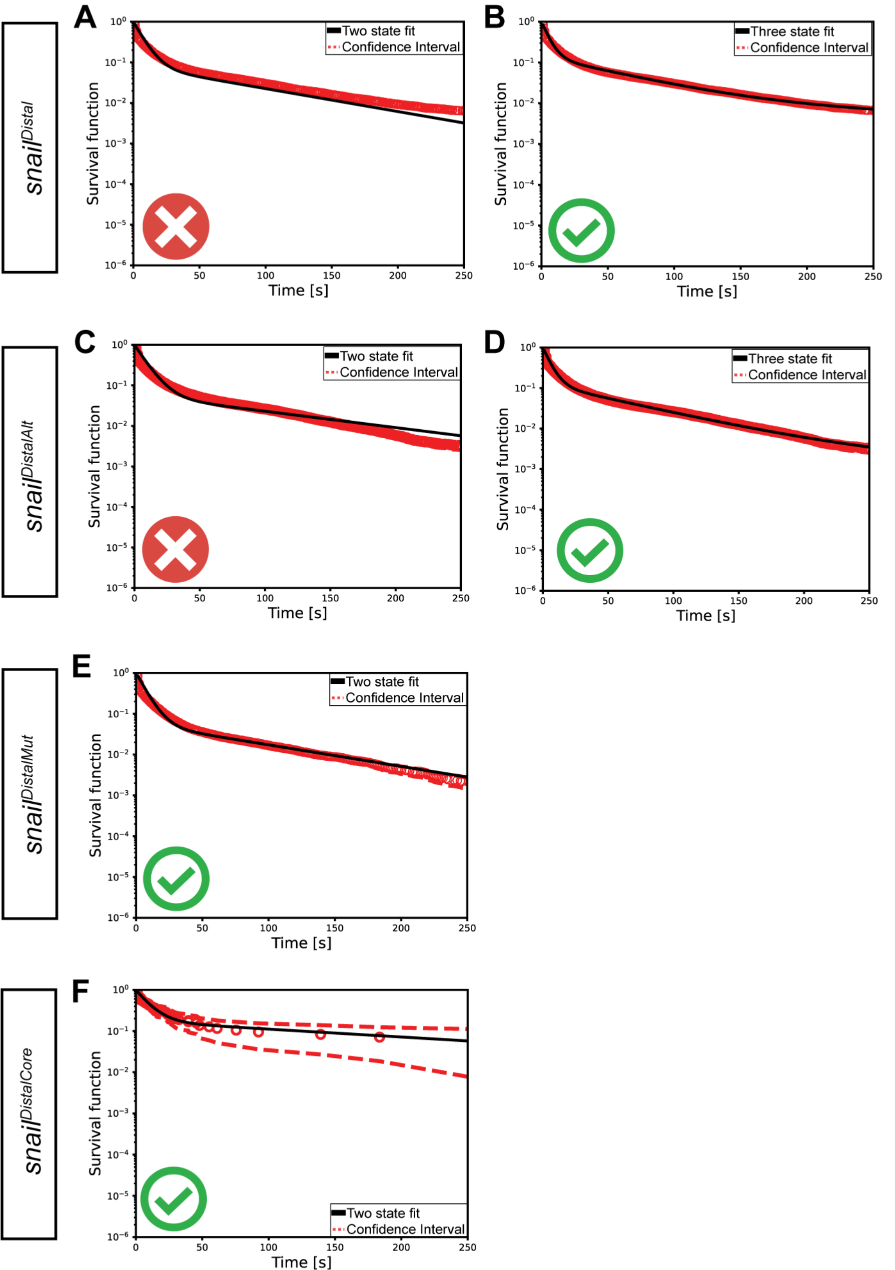

A
B
C
D
E
F
snail^Distal
snail^DistalAlt
snail^DistalMut
snail^DistalCore
Two state fit
Three state fit
Confidence Interval
Survival function
Time [s]

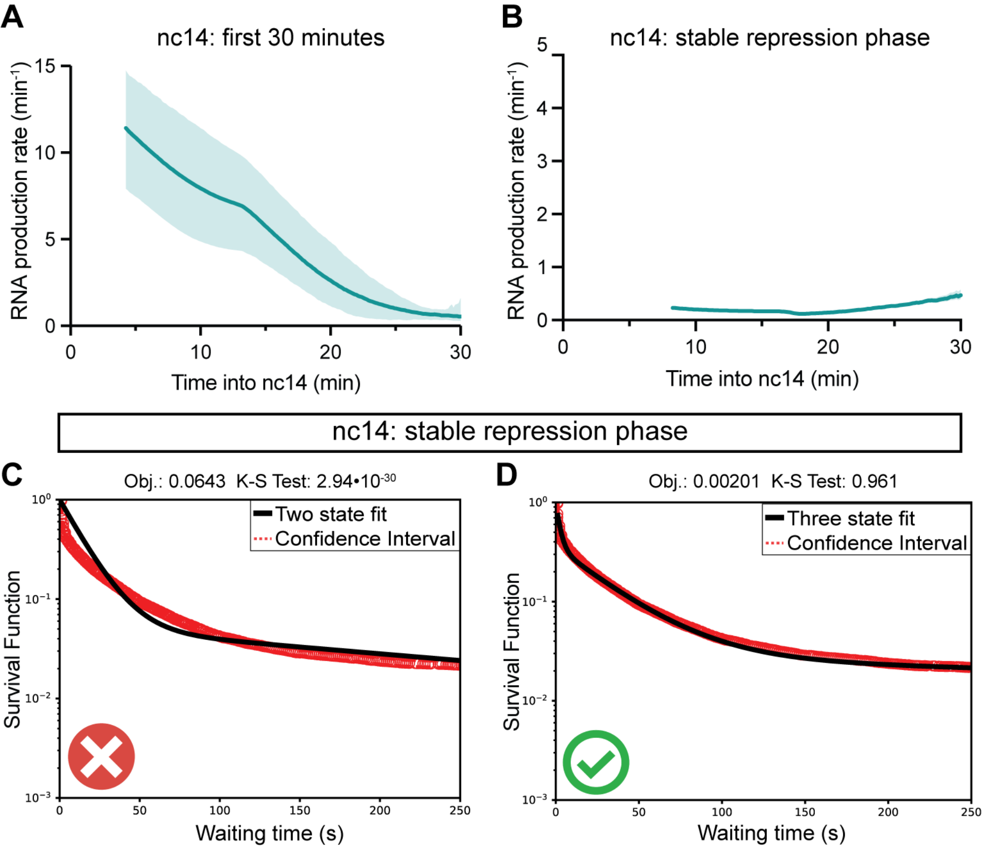

A
nc14: first 30 minutes
RNA production rate (min-1)
Time into nc14 (min)
B
nc14: stable repression phase
RNA production rate (min-1)
Time into nc14 (min)
nc14: stable repression phase
C
Obj.: 0.0643 K-S Test: 2.94•10-30
Two state fit
Confidence Interval
Survival Function
Waiting time (s)
D
Obj.: 0.00201 K-S Test: 0.961
Three state fit
Confidence Interval
Survival Function
Waiting time (s)

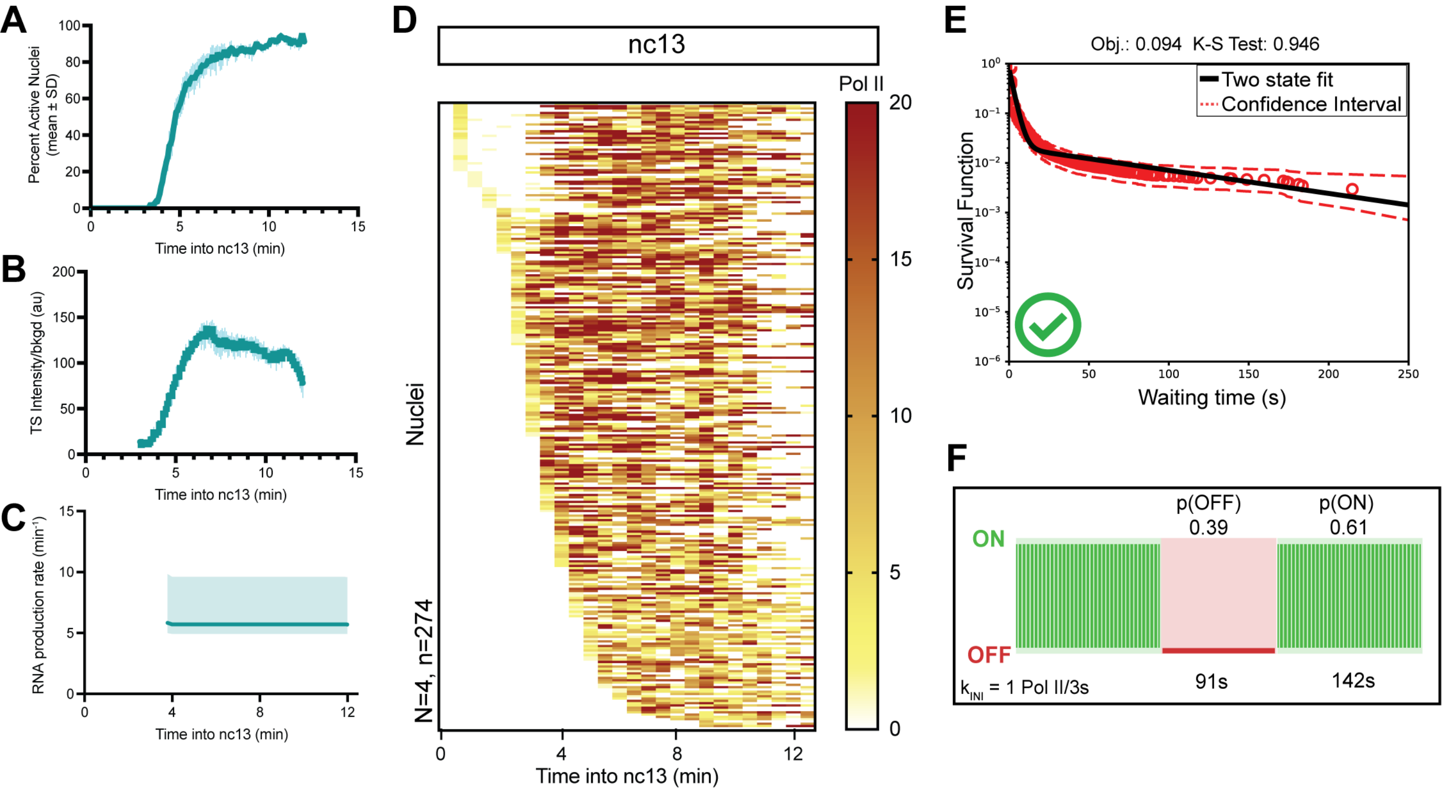

A
Percent Active Nuclei (mean ± SD)
Time into nc13 (min)
B
TS Intensity/bkgd (au)
Time into nc13 (min)
C
RNA production rate (min⁻¹)
Time into nc13 (min)
D
nc13
Pol II
Nuclei
N=4, n=274
Time into nc13 (min)
E
Obj.: 0.094 K-S Test: 0.946
Two state fit
Confidence Interval
Survival Function
Waiting time (s)
F
p(OFF)
0.39
p(ON)
0.61
ON
OFF
k_INI = 1 Pol II/3s
91s
142s

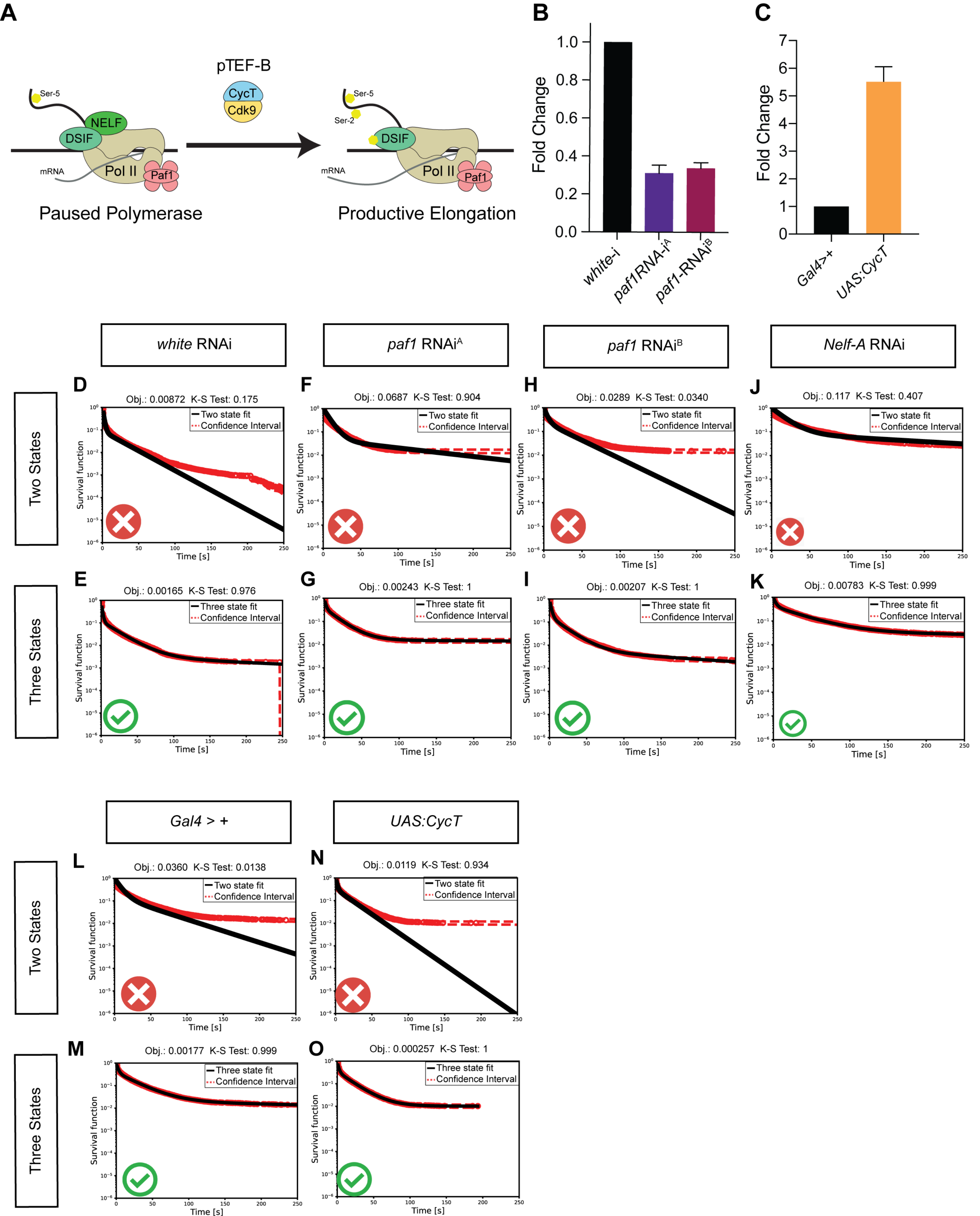

A
pTEF-B
CycT
Cdk9
Ser-5
Ser-2
NELF
DSIF
Pol II
Paf1
mRNA
Paused Polymerase
Productive Elongation
B
Fold Change
1.0
0.8
0.6
0.4
0.2
0.0
white-i
paf1RNA-i^A
paf1-RNAi^B
C
Fold Change
7
6
5
4
3
2
1
0
Gal4>+
UAS:CycT
white RNAi
paf1 RNAi^A
paf1 RNAi^B
Nelf-A RNAi
Two States
Three States
D
Obj.: 0.00872 K-S Test: 0.175
E
Obj.: 0.00165 K-S Test: 0.976
F
Obj.: 0.0687 K-S Test: 0.904
G
Obj.: 0.00243 K-S Test: 1
H
Obj.: 0.0289 K-S Test: 0.0340
I
Obj.: 0.00207 K-S Test: 1
J
Obj.: 0.117 K-S Test: 0.407
K
Obj.: 0.00783 K-S Test: 0.999
Two state fit
Three state fit
Confidence Interval
Survival function
Time [s]
Gal4 > +
UAS:CycT
L
Obj.: 0.0360 K-S Test: 0.0138
M
Obj.: 0.00177 K-S Test: 0.999
N
Obj.: 0.0119 K-S Test: 0.934
O
Obj.: 0.000257 K-S Test: 1

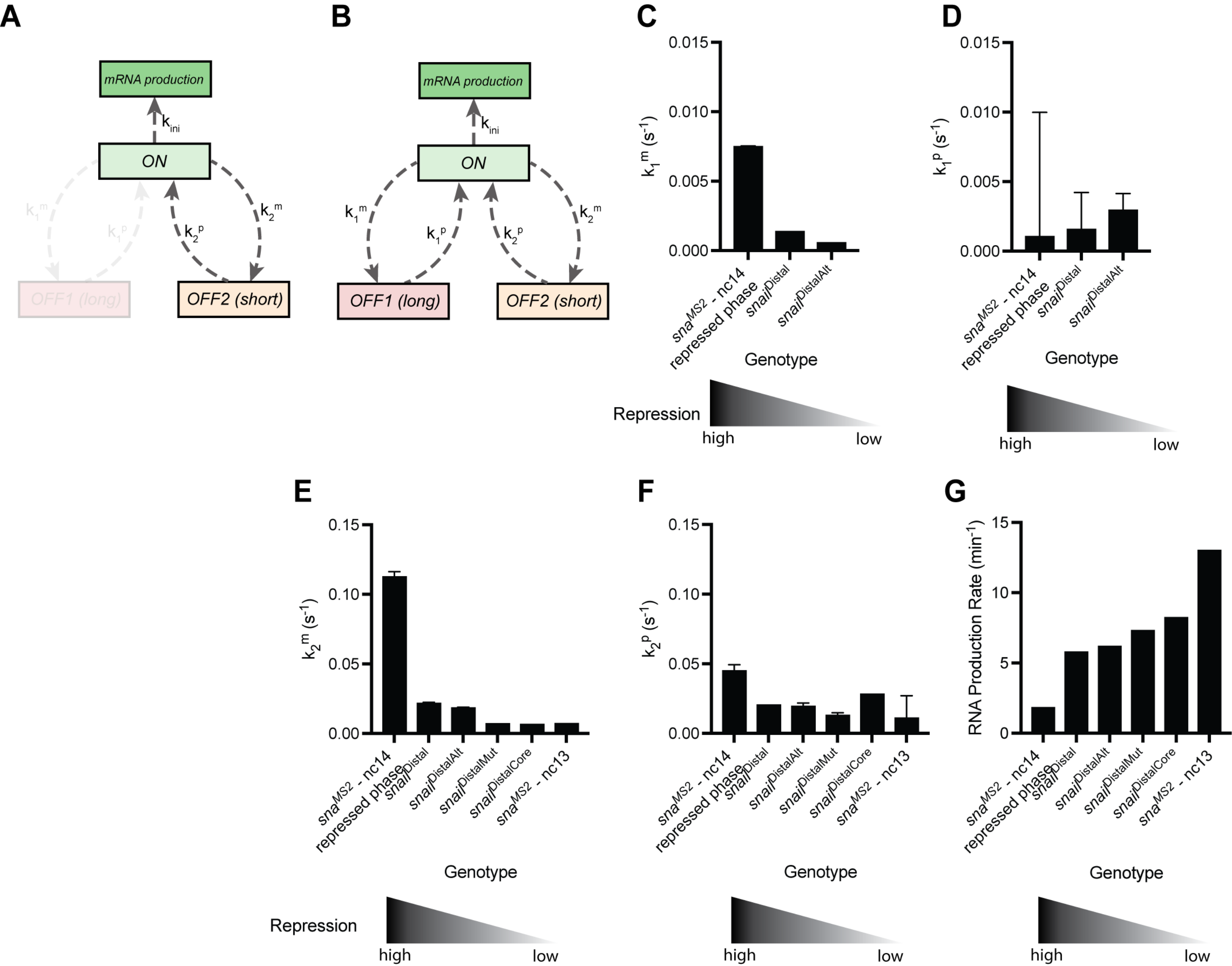
A
B
C
D
E
F
G
mRNA production
$k_{ini}$
ON
$k_1^m$
$k_1^p$
$k_2^p$
$k_2^m$
OFF1 (long)
OFF2 (short)
$k_1^m$ ($s^{-1}$)
$k_1^p$ ($s^{-1}$)
$k_2^m$ ($s^{-1}$)
$k_2^p$ ($s^{-1}$)
RNA Production Rate ($min^{-1}$)
0.015
0.010
0.005
0.000
0.15
0.10
0.05
0.00
15
10
5
0
$sna^{MS2}$ - nc14 repressed phase
$snail^{Distal}$
$snail^{DistalAlt}$
$snail^{DistalMut}$
$snail^{DistalCore}$
$sna^{MS2}$ - nc13
Genotype
Repression
high
low

**Supplemental Table 1:** Kinetic Parameters

| Genotype | 2 States Model | | | | | | | | | |
|---|---|---|---|---|---|---|---|---|---|---|
| | **k1+** | **k1-** | **k2** | **T(OFF) (s)** | **T(On) (s)** | **Pol II Initiaiton (s/event)** | **p(OFF)** | **p(ON)** | **Objective function** | **Kolmogorov-Smirnov** |
| ***sna* MS2 - active phase (nc13)** | **0.011** | **0.007** | **0.350** | **91** | **142** | **2.85** | **0.392** | **0.608** | **0.095432177** | **0.985401194** |
| Minimum | 0.011 | 0.005 | 0.330 | 91 | 192 | 3.03 | 0.161 | 0.608 | | |
| Maximum | 0.027 | 0.007 | 0.350 | 37 | 142 | 2.85 | 0.392 | 0.839 | | |

| Genotype | 3 States Model | | | | | | | | | | | | | |
|---|---|---|---|---|---|---|---|---|---|---|---|---|---|---|
| | **k1+** | **k1-** | **k2+** | **k2-** | **k3** | **T(OFF1) (s)** | **T(OFF2) (s)** | **T(On) (s)** | **Pol II Initiaiton (s/event)** | **p(OFF1)** | **p(OFF2)** | **p(ON)** | **Objective function** | **Kolmogorov-Smirnov** |
| **snaMS2 - repressed phase (nc14)** | **0.001** | **0.007** | **0.045** | **0.112** | **0.271** | **956** | **22** | **8** | **3.68** | **0.671** | **0.236** | **0.094** | **0.002011144** | **0.961125388** |
| Minimum | 0.001 | 0.007 | 0.045 | 0.112 | 0.271 | 956 | 22 | 8 | 3.68 | 0.184 | 0.236 | 0.094 | | |
| Maximum | 0.010 | 0.008 | 0.049 | 0.116 | 0.284 | 100 | 20 | 8 | 3.52 | 0.671 | 0.573 | 0.243 | | |

| Genotype | 3 States Model | | | | | | | | | | | | | |
|---|---|---|---|---|---|---|---|---|---|---|---|---|---|---|
| | **k1+** | **k1-** | **k2+** | **k2-** | **k3** | **T(OFF1) (s)** | **T(OFF2) (s)** | **T(On) (s)** | **Pol II Initiaiton (s/event)** | **p(OFF1)** | **p(OFF2)** | **p(ON)** | **Objective function** | **Kolmogorov-Smirnov** |
| ***snailDistal*** | **0.002** | **0.001** | **0.020** | **0.022** | **0.171** | **640** | **49** | **44** | **5.84** | **0.299** | **0.361** | **0.340** | **0.005894326** | **0.059164976** |
| Minimum | 0.002 | 0.000 | 0.020 | 0.022 | 0.171 | 237 | 49 | 44 | 5.75 | 0.029 | 0.361 | 0.340 | | |
| Maximum | 0.004 | 0.001 | 0.020 | 0.022 | 0.174 | 640 | 49 | 44 | 5.84 | 0.299 | 0.510 | 0.462 | | |
| ***snailDistalAlt*** | **0.003** | **0.001** | **0.019** | **0.018** | **0.160** | **342** | **52** | **53** | **6.23** | **0.089** | **0.443** | **0.468** | | |
| Minimum | 0.003 | 0.000 | 0.019 | 0.018 | 0.160 | 241 | 46 | 52 | 6.08 | 0.042 | 0.443 | 0.468 | | |
| Maximum | 0.004 | 0.001 | 0.022 | 0.019 | 0.165 | 342 | 52 | 53 | 6.23 | 0.089 | 0.446 | 0.512 | | |

| Genotype | 2 States Model | | | | | | | | | |
|---|---|---|---|---|---|---|---|---|---|---|
| | **k1+** | **k1-** | **k2** | **T(OFF) (s)** | **T(On) (s)** | **Pol II Initiaiton (s/event)** | **p(OFF)** | **p(ON)** | **Objective function** | **Kolmogorov-Smirnov** |
| ***snailDistalMut*** | **0.013** | **0.007** | **0.135** | **78** | **145** | **7.40** | **0.351** | **0.649** | **0.040272118** | |
| Minimum | 0.013 | 0.006 | 0.134 | 68 | 145 | 7.40 | 0.298 | 0.649 | | |
| Maximum | 0.015 | 0.007 | 0.135 | 78 | 159 | 7.45 | 0.351 | 0.702 | | |
| ***snailDistalCore*** | **0.028** | **0.006** | **0.124** | **36** | **154** | **8.10** | **0.188** | **0.812** | **0.028555408** | |
| Minimum | 0.028 | 0.006 | 0.124 | 36 | 154 | 8.10 | 0.188 | 0.812 | | |
| Maximum | 0.028 | 0.006 | 0.124 | 36 | 154 | 8.10 | 0.188 | 0.812 | | |

**Supplemental Table 2:** Fly lines associated with Pimmett et al. (2024)

| Line | In Text Reference | Reference |
|---|---|---|
| *; ; nos>* MCP-eGFP, His2A-mRFP | | Gift from T.Fukaya |
| ; *Mat-alpha* :GAL4/CyO; *nos* :GAL4, *nos* >MCP-eGFP, His2A-RFP/*nos* :GAL4, nos>MCP-eGFP, His2A-RFP | | Lagha lab |
| ; *snailMS2-3xP3-dsRed/snailMS2-3xP3-dsRed ;* | *snaMS2* | This paper |
| ; *sna ΔATG /* CyO, *hb* >lacZ ; | *snaΔATG* | This paper |
| *sogMS2/sogMS2 ; ;* | *sogMS2* | Whitney et al., Development 2022 |
| ; *snailLlama/snailLlama* ; | *SnailLlama* | This paper |
| *yw; P{w[+mC] = EGFP-STOP-bcd} ;* | *bcd>* GFP | Bothma et al., Cell 2018 |
| *w* ; *P{w[+mC]=His2Av-mRFP}/ CyO ;* | His2A-RFP | |
| *w1118* ; *P{GD9782}v20876* ; | Paf1 RNAi-A | VDRC 20876 |
| ; ; *P{KK100080}VIE-260B* | Paf1 RNAi-B | VDRC 108826 |
| ; ; *P{UASp-CycT.H}* | UAS:CycT | Hunt et al., Genome Biology 2024 |
| ; ; *P{y[+t7.7] v[+t1.8]=TRiP.HMS00686}attP2* | Nelf-A RNAi | BDSC 32897 |
| ; *PBac{sna-MS2-y}* ; | *snaWT* BAC | Bothma et al., eLife 2015 |
| ; *PBac{snaΔprimary-MS2-y}* ; | *snaΔPROX* BAC | Bothma et al., eLife 2015 |
| ; *PBac{snaΔshadow-MS2-y}* ; | *snaΔDIST* BAC | Bothma et al., eLife 2015 |
| ; ; PBPhi(*snaDistal-24xMS2-y)* (VK33) | *snaDistal* | Dufourt et al., Nature Communications 2018 |
| ; ; PBPhi(*snaDistalAlt-24xMS2-y)* (VK33) | *snaDistalAlt* | This paper |
| ; ; PBPhi(*snaDistalMut-24xMS2-y)* (VK33) | *snaDistalMut* | This paper |
| ; ; PBPhi(*snaDistalCore-24xMS2-y)* (VK33) | *snaDistalCore* | Ferraro et al., Current Biology 2016 |

**Supplemental Table 3:** guide RNA sequences for generation of CRISPR alleles

| Target | Sequence |
|---|---|
| *snail – MS2* | CGACATATGAATCCCTTAGCAGG |
| *snail – Llama* | CGACATATGAATCCCTTAGCAGG, CCCCATGAACGAAGAGTACTAGG |
| *snail – ΔATG* (guide) | GTAGTGACCCATTGAATTCGTGG |
| *snail – ΔATG* (ssODN)<br>**guide sequence**<br>mutations<br>EcoRI site for screening | TCGATCAGTACCGGAAACTAAAACTTAATCACACACACATCAAAAATGGCCGCC AACTACAAAAGCTGCCCGCTAAA**GTAGTGACCCATTGAATTCGTGG**AGGAGC GTCTGCCACAAACGGAGGCCTTGGCCCTGACCAAGGACTCACAGTTTGCCCA GGATCAGCCGCAGGATCTATCCCTGAAACGGGGTCGCGACGA |
| *White - coffee* | ATACCATTCCTGCTCTTTGG |

**Supplemental Table 5:** single molecule FISH probes

*snail* endogenous smFISH probes

| Probe | Sequence | Probe Type |
|---|---|---|
| snail_1 | tctcaacgagagctgaggtg | smFISH |
| snail_2 | gagtatagagcggtggttgc | smFISH |
| snail_3 | tgggtaaatcgggagatcgg | smFISH |
| snail_4 | agttttagtttccggtactg | smFISH |
| snail_5 | ccatttttgatgtgtgtgtg | smFISH |
| snail_6 | ttagcgggcagcttttgtag | smFISH |
| snail_7 | ctcctccacgaagacaatgg | smFISH |
| snail_8 | caaactgtgagtccttggtc | smFISH |
| snail_9 | cgtttcagggatagatcctg | smFISH |
| snail_10 | tgctgataatcctgggtctc | smFISH |
| snail_11 | acatagtcacgtttcggttc | smFISH |
| snail_12 | ccggtgtttttgaaaggttc | smFISH |
| snail_13 | agttggagctagagctggag | smFISH |
| snail_14 | tagtccacgcatatggattt | smFISH |
| snail_15 | gattaatcgtggtgggggtg | smFISH |
| snail_16 | atcacaaaggcggactggaa | smFISH |
| snail_17 | cagagatcggattgcaaccg | smFISH |
| snail_18 | atctgctggtagctgtagac | smFISH |
| snail_19 | aaccggtttccagatcggat | smFISH |
| snail_20 | actgaaagatcctctggctc | smFISH |
| snail_21 | cggcagtgggatgtcatttc | smFISH |
| snail_22 | gcctcatcgaaaaggtggaa | smFISH |
| snail_23 | tgtaggagtatcccgatgag | smFISH |
| snail_24 | catgattggcggcaacactc | smFISH |
| snail_25 | gcacttgaagcggtagtttt | smFISH |
| snail_26 | atcgaggtggagtacatctt | smFISH |
| snail_27 | aactgacggtgcttggacag | smFISH |
| snail_28 | ttcttctcctgattacactc | smFISH |
| snail_29 | aatggtggtgtacagctttc | smFISH |
| snail_30 | gtgcggatgtgcatcttcag | smFISH |
| snail_31 | caaatggggcacttgcaggg | smFISH |
| snail_32 | agggtcgagagaaggccttg | smFISH |
| snail_33 | aaaggcttctctccagtgtg | smFISH |
| snail_34 | caaaggatcgtgggcagtcg | smFISH |
| snail_35 | gatgagctcgcaggttcgag | smFISH |
| snail_36 | tacttcttgacgtccacgtg | smFISH |
| snail_37 | gaaagatttgtggcacacct | smFISH |
| snail_38 | tgcttgttcaggagcgacat | smFISH |
| snail_39 | tagtgatggtgcagttggag | smFISH |
| snail_40 | atatgtcgagaatcctacgc | smFISH |
| snail_41 | taattgtgtcctgctaaggg | smFISH |
| snail_42 | gcggaatgtgagtttgctta | smFISH |
| snail_43 | attgtctgtttgtttggtct | smFISH |
| snail_44 | gcaccaaaaccgaatcgact | smFISH |
| snail_45 | atgctgcgtgtgacaatgag | smFISH |
| snail_46 | acagttggcttaacagtact | smFISH |
| snail_47 | ttcttctctttaagctagga | smFISH |

*MS2* smiFISH probes

| Probe | Sequence | Probe Type |
|---|---|---|
| MS2_1 | GATCGTCGTCGTTTGAAGATTCGACCTGG | smiFISH |
| MS2_2 | CGGCTGATGCTCGTGCTTTCTTGGCA | smiFISH |
| MS2_3 | CGTAGGATCTGATGAACCCTGGAATACTG[illegible] | smiFISH |

*yellow* smFISH probes

| Probe | Sequence | Probe Type |
|---|---|---|
| yellow_1 | atcagggtcacaagaatcca | smFISH |
| yellow_2 | actatatcgctcctgaagtt | smFISH |
| yellow_3 | tttagtcgggtattcgggaa | smFISH |
| yellow_4 | tataatctccactagccaga | smFISH |
| yellow_5 | ttcgactccaacaggtagag | smFISH |
| yellow_6 | gtgacgaataaccgattgcc | smFISH |
| yellow_7 | aaactgcggtccatgtttat | smFISH |
| yellow_8 | gccaatctggatacggaatt | smFISH |
| yellow_9 | caatctccagctgtatttga | smFISH |
| yellow_10 | gtaggcagtggtaatactgt | smFISH |
| yellow_11 | ccacactcatccactttaat | smFISH |
| yellow_12 | cacggattagtggtggtatt | smFISH |
| yellow_13 | gtatccgtggtcaagtcaaa | smFISH |
| yellow_14 | tagctcgtatctccgaattc | smFISH |
| yellow_15 | gtatttggatttgtgtccac | smFISH |
| yellow_16 | cacggcaatgttagctatga | smFISH |
| yellow_17 | atcatcgcaatttttgccta | smFISH |
| yellow_18 | tatcccaattcatcggcaaa | smFISH |
| yellow_19 | cccaggagtaagcaatcaag | smFISH |
| yellow_20 | agaatctccaggacttgttc | smFISH |
| yellow_21 | cctcaatggatcggggaaaa | smFISH |
| yellow_22 | cccattggaagttaatacca | smFISH |
| yellow_23 | ataccaaatataccctcctc | smFISH |
| yellow_24 | cgatcgaatgggcgaaaggg | smFISH |
| yellow_25 | agtacagggtacgataacca | smFISH |
| yellow_26 | cgatgacttgctaacggact | smFISH |
| yellow_27 | aaaatcctcgtggatacggc | smFISH |
| yellow_28 | catgatagctatcttccgtc | smFISH |
| yellow_29 | ccgttcatctaaggcaacaa | smFISH |
| yellow_30 | cacgtgaagtggtatgggag | smFISH |
| yellow_31 | acagctcaattccatcatcg | smFISH |
| yellow_32 | gagtacggcattgatgagtg | smFISH |
| yellow_33 | ccacaatgccatgaaattgc | smFISH |
| yellow_34 | ttttcacatcggccggaaaa | smFISH |
| yellow_35 | acccaaacgtttttgttctc | smFISH |
| yellow_36 | gcaagaaaacgggcatccta | smFISH |
| yellow_37 | aagggagccgtgtaaattcg | smFISH |
| yellow_38 | aggcgttattcctcaaatca | smFISH |
| yellow_39 | acggcttgttttggtattga | smFISH |
| yellow_40 | atataacggtggacccattg | smFISH |
| yellow_41 | tttctgtggcaagacaggac | smFISH |
| yellow_42 | cgggcaaataagtgcgactt | smFISH |
| yellow_43 | tggagactacattgcctgaa | smFISH |
| yellow_44 | ggacccacagaatttgtaga | smFISH |
| yellow_45 | ccgttgtgctggttgaaaat | smFISH |
| yellow_46 | gaccacttgtctcgtaattt | smFISH |
| yellow_47 | gggttgatgggtgggaaata | smFISH |
| yellow_48 | aaccttgatgctgatgatgc | smFISH |

**Supplemental Table 6**: qPCR Primers

| Name | Sequence |
|---|---|
| paf1_F | CACCGCTTCGTGCAGTACAA |
| paf1_R | CCAAATCGTGTTCCGTCAGC |
| CycT_F | CCGGCCCGTCTGAAGTCTA |
| CycT_R | CCTTGCTGTTAGCTGTCCGAT |
| Rpl13_F | AGCGGCATGTGAAGACCTG |
| Rpl13_R | AAGACGGCCTTAGCCTTCTTG |
